# Two-way feedback between chromatin compaction and histone modification state explains *S. cerevisiae* heterochromatin bistability

**DOI:** 10.1101/2023.08.12.552948

**Authors:** Ander Movilla Miangolarra, Daniel S Saxton, Zhi Yan, Jasper Rine, Martin Howard

## Abstract

Compact chromatin is closely linked with gene silencing in part by sterically masking access to promoters, inhibiting transcription factor binding and preventing polymerase from efficiently transcribing a gene. Here, we propose a broader view: chromatin compaction can be both a cause and a consequence of the histone modification state, and this tight bidirectional interaction can underpin bistable transcriptional states. To test this theory, we developed a mathematical model for the dynamics of the HMR locus in *S. cerevisiae*, that incorporates activating histone modifications, silencing proteins and a dynamic, acetylation-dependent, three-dimensional locus size. Chromatin compaction enhances silencer protein binding, which in turn feeds back to remove activating histone modifications, leading to further compaction. The bistable output of the model was in good agreement with prior quantitative data, including switching rates from expressed to silent states, and vice versa, and protein binding levels within the locus. We then tested the model by predicting changes in switching rates as the genetic length of the locus was increased, which were then experimentally verified. This bidirectional feedback between chromatin compaction and the histone modification state may be an important regulatory mechanism at many loci.

**Significance:** Chromatin is the complex formed by proteins, including histones, and DNA to form chromosomes. Specific chromatin structures and states are thought to be key factors regulating transcription. A common view proposes that histone modifications activate or inhibit transcription either via specific activation or inhibition of RNA polymerase binding/elongation at a locus, or by expanding/compacting the locus, thereby modulating its accessibility to many macromolecules. In this work, we elucidated a broader hypothesis that chromatin compaction may both inhibit transcription, and feedback via silencing proteins to remove histone modifications that further control chromatin compaction and correlate with gene activity. We developed a model incorporating these ideas and showed that it explains quantitative experimental data for a silent locus in budding yeast.

---

Although the genome sequence underpins much of the biology of an organism, cells with the same genotype can nevertheless have very different phenotypes, mediated through differential regulation of gene expression. Gene expression is in turn modulated by the multifaceted characteristics of chromatin.

For almost a century, chromatin has been broadly divided in euchromatin, open and generally associated with active genes, and heterochromatin, compact and silent (Passarge, 1979). Coherent with this vision, the three-dimensional arrangement of chromatin in the vicinity of a gene and the post translational modification (PTMs) of the nearby nucleosomes are thought to be major determinants of the strength of transcription. Nevertheless, the relationship between the three-dimensional structure and chromatin modifications has not been elucidated in most cases. In fact, there is lively debate regarding which is the actual transcriptional regulator: whether it is the local density of chromatin (Ou et al., 2017), histone PTMs present in the locus (Allshire & Madhani, 2018), or the identity of the DNA or nucleosome-bound proteins (Kueng et al., 2013).

A common view is that if chromatin is highly compacted, the DNA in this region will be inaccessible and transcriptionally silent (Klemm et al., 2019; Ou et al., 2017). Conversely, if the chromatin is less densely packed, then transcription factors can bind, which will be followed by RNA polymerases (Preissl et al., 2022). However, accessibility and transcription are not always correlated (DeVeale et al., 2022; Kiani et al., 2022).

Key regulators of chromatin compaction are reader-writer enzymes that bind to nucleosomes only if certain histones PTMs are present and which catalyse the spread of that same histone PTM to nearby nucleosomes. For example, PRC2 is activated by H3K27me3 to add further H3K27me3 modifications nearby in a read-write feedback, driving chromatin compaction in vitro (Grau et al., 2021). Other reader-writer enzymes have also been suggested to be involved in chromatin compaction, such as the SIR complex in budding yeast (Ruault et al., 2021). However, there are alternative regulators of chromatin compaction. Significant evidence points towards acetylation of histone tails, especially H4, as controlling the physical conformation of chromatin (Collepardo-Guevara et al., 2015; Robinson et al., 2008).

These concepts are often presented in a sequential manner, where upstream effectors and downstream consequences are clearly distinguished. Here, however, we elucidated a more nuanced description, where the extent of chromatin compaction is controlled by the absence of histone PTMs, but where the absence of histone PTMs is itself influenced by the extent of chromatin compaction. Indeed, many reader-writer enzymes dimerise (e.g., PRC2) and bind to more than one nucleosome at the same time, which makes them very sensitive to the local physical configuration of chromatin. A sufficiently compacted locus would increase the binding of these enzymes and would thereby further influence the histone marks that the enzymes spread or remove. As the presence or absence of histone marks can modulate chromatin conformation, the physical state of the chromatin environment and the histone modification state are inextricably linked in a mutually reinforcing manner.

In this work, we developed this hypothesis via a mathematical model for the *S. cerevisiae* silenced mating-type locus *HMR*. This locus is constitutively silenced in budding yeast by the action of the Silent Information Regulator (SIR) proteins, which form the SIR complex. The binding of the SIR complex to the locus causes two major changes in the chromatin: deacetylation of H4K16 (H4K16 is mostly acetylated in yeast) and loss of H3K79 methylation (Rusche et al., 2003). Furthermore, *HMR* is flanked by two silencers (E and I), which are regulatory sites that recruit Sir proteins and which are essential for heterochromatin formation at the locus, as their deletion leads to loss of silencing. The genomic regions outside the locus and next to the silencers, including a tRNA gene, act as insulators, preventing the spread of heterochromatin beyond the boundaries of the locus (Donze et al., 1999). Moreover, silencing at *HMR* does not depend on the identity of the gene encoded by the locus, allowing empirical observation of the transcriptional state of the locus by substituting reporter genes for those genes usually resident at *HMR*.

Under certain conditions that weaken the action of the silencers, the silencing of the locus is rendered bistable, switching between silenced and expressed states. The transcriptional state can be inherited over multiple generations, depending on the strength of silencers, making it an ideal system to study the inheritance of epigenetic information to daughter cells. Capturing its dynamics quantitatively requires an accurate mathematical model, with detailed experimental data available as a strict benchmark for our theory. Here, model outcomes for the *HMR* locus were successfully tested against both prior experimental data and the results of new experiments specifically designed to test our hypothesis. Overall, our model provided a faithful representation of the biochemical processes involved in epigenetic silencing, enabling us to develop a coherent theory for the feedback between the histone PTM state and chromatin conformation in three-dimensional space.

Previous modelling studies have focused either on a coarse-grained description of large-scale chromatin or on a detailed picture of the regulation at a particular locus. In the coarse-grained description a similar bidirectional interaction between epigenetic state and chromatin folding is considered, but the molecular mechanisms modelled are often abstract and with limited detail, making quantitative comparison with experimental data difficult (Jost & Vaillant, 2018; Michieletto et al., 2018; Michieletto et al., 2016; Owen et al., 2022; Pease et al., 2021; Sandholtz et al., 2020). In parallel, single-locus studies have provided much insight into the regulation of certain genetic regions, capturing their dynamics quantitatively, but they have not incorporated this link between chromatin and the histone PTM state (Angel et al., 2011; Berry et al., 2017; Nickels et al., 2021; Sneppen & Dodd, 2015). In this work, we drew inspiration from both approaches to produce a model that included this important link and, at the same time, kept a detailed description of the molecular mechanisms regulating *HMR* heterochromatin formation, enabling an extensive quantitative comparison with experimental data. Indeed, the *HMR* locus is one of a small number of loci from any organism where sufficient detailed molecular data exists to make such an in-depth comparison.

## Results

### Underlying processes and timescales

Intensive study of the budding yeast *HMR* locus has identified proteins, enzymes, histone PTMs and regulatory sites involved in silencing/activation (Kueng et al., 2013; Rusche et al., 2003). Sas2 acetylates H4K16 and Dot1 methylates H3K79, especially if the H3 histone substrate is bound to an H4 histone carrying the H4K16ac mark (Valencia-Sánchez et al., 2021). These two PTMs are associated with active loci. With respect to silencing, the silencers delimiting the locus have significant affinity for proteins such as ORC and Rap1 that recruit Sir1 which, in turn, recruits the rest of the SIR complex. If the nucleosome is unmodified, the SIR complex (Sir2-3-4) can bind to it, but in presence of the H4K16ac or H3K79me marks, the Sir complex affinity is greatly reduced (Rusche et al., 2003). Spreading of the SIR complex is then mediated by the read-write action of Sir2, a histone deacetylase whose preferred substrate for the enzymatic reaction are histones acetylated at H4K16, thus increasing the number of binding sites for other Sir proteins by means of its catalytic activity. Notably, binding of the SIR complex to nucleosomes occurs in a highly cooperative manner as its affinity increases roughly 50-fold when it binds to two unmodified nucleosomes (Behrouzi et al., 2016). Importantly, *sir1Δ* strains are bistable with respect to its transcriptional state at HMR and are able to maintain such a state for multiple generations (Pillus & Rine, 1989). Thus, in a *sir1Δ* mutant, HMR is found either in an active state (with high levels of acetylation) or in a silenced state (with low levels of H4K16ac and H3K79me3, but with high levels of the SIR complex bound), switching between the two typically on a timescale of tens of generations (Saxton & Rine, 2022).

Given the variety of processes involved in the silencing of the locus, from protein binding to DNA replication, estimating the timescales for these dynamical processes is important for a comprehensive and quantitative understanding of silencing. The fastest process is Sir protein binding and unbinding, on the order of tens of seconds (Behrouzi et al., 2016). Acetylation and deacetylation of histone H4 tails occurs at an intermediate timescale, around 10 minutes (Waterborg, 2001). Finally, since there is no known demethylase in yeast for H3K79, the only way to lower the number of H3K79me3 marked histones is by dilution at replication (Goodnight & Rine, 2020). Hence, the timescale of H3K79 methylation is on the order of the duration of the cell cycle, i.e., approximately 2 hours. The model we developed, described in the next section, reflects these experimental findings.

### Modelling histone PTM dynamics

Here we summarise the model constituents and fundamental dynamics (depicted schematically in Fig. 1A; for full details and parameter values, see *SI Appendix, Sections 1 and 2*).

**Figure 1.**
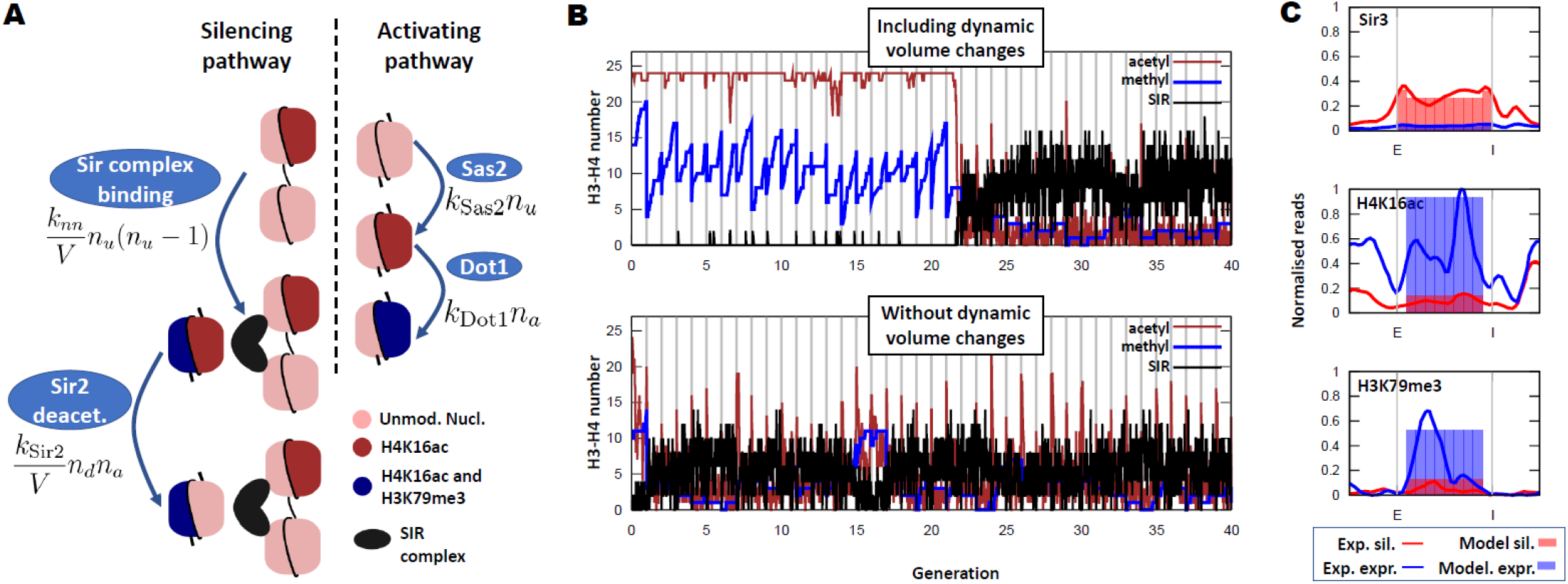
A model with dynamic volume changes can reproduce the bistable dynamics of HMR in the sir1Δ mutant. A) Schematic of the model. The SIR complex binds by dimerisation to two unmodified H3-H4 pairs and Sir2 deacetylates H4 histones (third nucleosome in the silencing pathway, left column). Sas2 acetylates H4 histones and Dot1 methylates acetylated H3-H4 histone pairs. k_nn_ is the reaction constant for the SIR complex binding to two unmodified nucleosomes, n_d_ is the number of H3-H4 histone pairs dimerised by the SIR complex, and k_Sir2_ the reaction constant for Sir2 deacetylation of H4K16. The rest of the symbols are defined in the main text. B) Example output of a simulation for the sir1Δ mutant. The full model (including volume changes) shows a robust bistable behaviour that persists over generations (similar to the phenotype found in experiments), while the same model with a fixed locus volume is neither bistable nor robust. C) The occupancy fraction of Sir proteins and histone PTMs are compared to experimental ChIP profiles for each of the stable states of the model (silenced [sil.] and expressed [expr.]). Experimental data obtained from Ref. (Saxton & Rine, 2022). ChIP profiles were normalised to their highest peak across the locus, in order to enable quantitative comparison with occupancy fraction.

Given that a nucleosome is formed by a histone octamer (two pairs of H2A-H2B histones and another two pairs of H3-H4 histones), we modelled the locus as a set of H3-H4 pairs which can be in five states: Unmodified (U), Unmodified and Sir bound (Ub), Acetylated (H4K16ac, A), Methylated (H3K79me3, M) and Acetylated and Methylated (AM). We assumed that Sir proteins can bind only to unmodified H3-H4 pairs by a dimerisation process, binding two H3-H4 histone pairs at a time, given that individually they have a very low affinity (Behrouzi et al., 2016). To reflect a compact and dynamic arrangement of the locus, the model does not include the relative positions of the nucleosomes, it treats the locus as well-mixed, where contacts between any two H3-H4 pairs are all equally likely. For simplicity, and given the insulator role of the boundaries of the HMR locus (Donze et al., 1999), we considered the locus isolated from the rest of the genome and we do not model the rest of the chromosome or any other genetic region.

The two silencers were modelled as preferred binding sites for the Sir complex, which can be thought of as H3-H4 pairs fixed in the U state but with a higher affinity for Sir proteins. Sir proteins can also dimerise between the two silencers or between a silencer and any unmodified H3-H4 pair. In a one-dimensional representation of the locus, a silencer would stand at each end of the locus and the nucleosomes would be stacked in between, implying that it would be easier for Sir proteins to dimerise between nearest neighbours. Nevertheless, this one-dimensional configuration of nucleosomes is not an accurate depiction of reality and the locus is likely to adopt a more dynamic and disordered configuration (Farr et al., 2021). To reflect this dynamic configuration, we allowed Sir proteins to dimerise between any pair of nucleosome and silencer; but to account for the longer relative length (in the one-dimensional picture) between silencers, and between silencers and nucleosomes (on average), we weighted the dimerization probabilities by a factor determined by the genetic length between them. Given that the average distance between any two nucleosomes in the locus is roughly 1/3 of the length of the locus, and that the average distance between a silencer and any nucleosome is 1/2 of the length of the locus, a weight of 2/3 was applied to the nucleosome-silencer binding constant. Analogously, a factor of 1/3 was applied to the silencer-silencer binding constant (see *SI Appendix Section 1*). Therefore, we accounted for longer genetic distances while still allowing for long-range binding, approximately reflecting the dynamic and folded architecture of the locus (Valenzuela et al., 2008). In cells with weakened silencers, we reduced the effective binding constant of the Sir complex to silencers, e.g., in the *sir1Δ* strain we reduced the binding affinity of silencers to almost that of nucleosomes.

The dynamics of the histone PTMs were modelled mathematically with a reaction rate proportional to the substrate amount, i.e., the rate of H4K16 acetylation by Sas2 is *k*_*Sas*2_*n*_*u*_, where n_u_ is the number of unmodified (and not SIR-bound) H3-H4 pairs in the locus at any point in time and *k*_*Sas*2_ is a constant reflecting the activity and nuclear concentration of Sas2. Similarly, Dot1 methylation of H3K79 is given by the rate *k*_*Dot*1_*n*_*ac*_, since the most efficient substrate of Dot1 are acetylated H3-H4 pairs (Valencia-Sánchez et al., 2021). The Sir2 deacetylation rate depends on the product of the amount of SIR complex at the locus and the acetylated H3-H4 pairs (see Fig. 1A for the mathematical form). We also considered that Sas2 and Sir2 can acetylate and deacetylate (respectively) H3-H4 pairs methylated at H3K79, with analogously constructed reaction rates. The acetylation and methylation processes comprise the activating pathway, while Sir complex binding and Sir2 deacetylation are defined as the silencing pathway (see Fig. 1A for a diagram of these PTMs and their rates). Finally, at the end of the cell cycle, replication was modelled with every nucleosome having a 50% probability of being replaced by one which is fully acetylated and unmethylated, since, in *S. cerevisiae*, most of the H4 histones in the nucleoplasm are acetylated (Shia et al., 2005). The 50% probability of replacing a nucleosome at replication is supported experimentally in budding yeast (Schlissel & Rine, 2019).

Crucially, Sir complex binding to histones and Sir2 deacetylation, the two steps in the silencing pathway, rely on contacts between regions within the locus. Contacts within the locus will depend on the three-dimensional locus configuration, which, in a first approximation, could be modelled by quantifying the three-dimensional volume occupied by the locus: large if the locus is open (with few contacts within the locus), or small if the locus is in a compact configuration (allowing many more contacts). In terms of polymer physics, we estimated the volume occupied by the locus, *V*, as proportional to 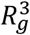, where *R*_*g*_ is the radius of gyration of the locus (Rubinstein et al., 2003). Therefore, the overall volume of the locus *V* modulates the rates of Sir binding and Sir2 deacetylation by changing the frequency of contacts between nucleosomes, see Fig. 1A. Moreover, there is evidence that H4K16 acetylation is critical in determining the physical conformation of chromatin (Collepardo-Guevara et al., 2015; Robinson et al., 2008), due to H4 tail-DNA contacts [according to molecular dynamics simulations (Farr et al., 2021) and *in vitro* experiments (Wang et al., 2013)]. Thus, drawing inspiration from polymer physics we modelled the varying volume of the locus with the following scaling:

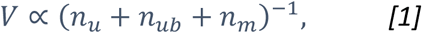

where *n*_*ub*_ is the number of unmodified and SIR-bound H3-H4 pairs, and *n*_*m*_ that of methylated H3-H4 pairs and *n*_*u*_ + *n*_*ub*_ + *n*_*m*_ is the total number of H3-H4 pairs that are not acetylated at H4K16. This scaling arises as a limiting case of the Flory picture for real polymers in a poor solvent (Rubinstein et al., 2003), when the coil is compact and the entropic contribution of the chain to the free energy can be neglected (see *SI Appendix, Section 3*). Thus, the overall locus volume is reduced in loci with many unmodified H4 tails while it expands if the levels of H4K16ac are high. Consistent with this picture, there are frequent contacts between the two silencers and the nucleosomes in the wild type, when the locus is fully silenced, but these contacts disappear in a *sir3*Δ mutant, where silencing is abolished (Valenzuela et al., 2008). Chromatin compaction has also been observed upon addition of catalytically active Sir proteins to chromatin *in vitro* presumably due to deacetylation (Johnson et al., 2009). Overall, these acetylation-dependent dynamics provide the link between the physical conformation of the locus and the histone PTMs.

The model was then implemented with a stochastic simulation algorithm (Gillespie, 1977). Importantly, when trying to recapitulate the features of the bistable *sir1Δ* mutant with the model we found that, if we assumed the volume was fixed and did not depend on the chromatin state, the bistability and robustness of the model became compromised. In Fig. 1B, both simulations were initialised in a highly expressed state (high acetylation and methylation) but, in the absence of dynamic volume changes, the active state was not stable. Generation of bistable behaviour required acetylation-dependent volume changes, arising from a fundamental difference between the processes that activate the locus and those that silence it. In the model, the silencing processes (Sir binding and Sir2 deacetylation) depend on contacts between nucleosomes within the locus, but the activating processes do not. As a result, an expanded locus would favour activation, whereas a compact one would promote silencing. The system could therefore become locked in a compact and silenced locus by the interaction between enzymes promoting loss of activating histone PTMs and the compact volume which enhances the binding of these enzymes. Stochastic silencing loss was rare but did occur (roughly every 100 generations) and when this happened the locus now locked itself into an expressed state, where the locus had expanded due to the higher acetylation making it difficult for the Sir complex to bind and reverse the changes. These two stable states were highly polarised, one of them having high Sir protein binding and low H4K16ac and H3K79me3, while the expressed state showed little protein binding and high levels of acetylation and methylation.

We also found quantitative agreement between the amount of Sir binding and histone PTMs in each of the two states (silenced and expressed), see Fig. 1C. In the simulations the locus is considered expressed if, at the end of the cell cycle but before DNA replication, half or more of the H3-H4 pairs are either acetylated or methylated, a reasonable threshold given recent experimental data (Wu et al., 2021). However, due to the bistable nature of the model, varying this threshold moderately did not produce significant changes to the model output. A quantitative comparison was then made with data from Ref. (Saxton & Rine, 2022), by renormalising the ChIP profiles to the highest peak, in an attempt to estimate the maximum occupancy, with good results. Note that the model did not include details of the position of the nucleosomes or any sequence-dependent effect other than the silencers, so the model results were not able to recapitulate the fine-scale spatial structure of chromatin. Overall, we found that the model effectively captured the known behaviour of the *HMR* locus.

### Switching rates in the *HMR* loci with different genetic lengths

Previously, two of us published the results of an experiment in which the genetic length of the *HMR* locus, and thus the number of nucleosomes between the silencers, was varied and the silencing establishment/loss switching rates were measured. If epigenetic memory of the transcriptional state of the locus is transferred to daughter cells by nucleosome inheritance, having fewer nucleosomes in the locus should make inheritance less stable, as there would be more variability in the fraction of nucleosomes inherited. However, this is not what was observed in Ref. (Saxton & Rine, 2019). There was no change in silencing loss rates and an increase, although statistically not significant, in the silencing establishment rates as the number of nucleosomes was reduced. Recapitulating these data is a strict test of the model, since it requires a fine balance between the volume of the locus, the number of nucleosomes and the scaling of the reaction rates locus wide.

With appropriate parameters, the model could recover the results obtained in that experiment, see Fig. 2A and B, if we assumed that the fluorescent protein measurements in the experiments are delayed by a few hours with respect to the chromatin state, due to subsequent transcription and translation. Consequently, in the statistics of the model, we did not consider switching events that are reversed in less than two generations, as such events would not be visible experimentally via a fluorescent protein assay. With these considerations, the model produced an important prediction: the silencing loss rate as a function of the number of nucleosomes in the locus remains approximately constant, while the silencing establishment was reduced substantially as the genetic locus size increases. It is worth noting that silencing loss rates are around 1%, a low rate that is enabled in the model by dynamic volume changes.

**Figure 2.**
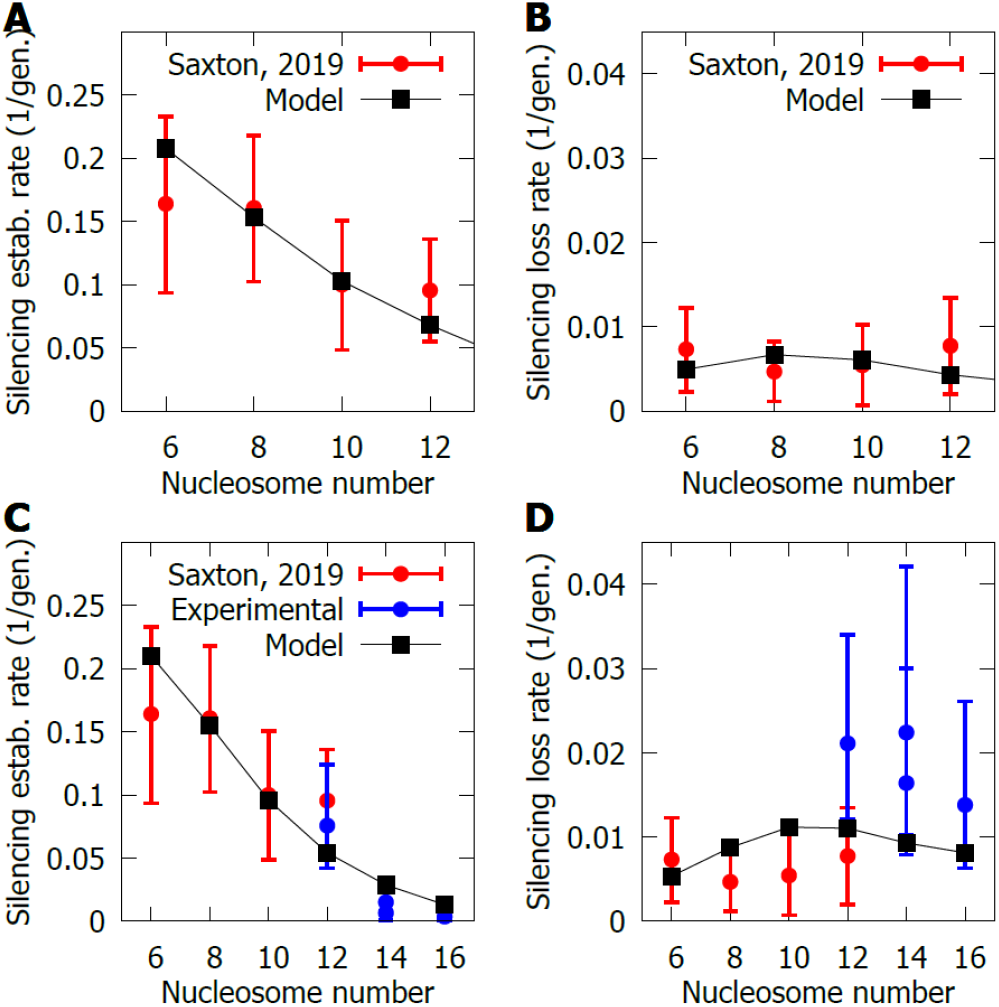
Switching rates as a function of nucleosome number. A,B) Silencing establishment and silencing loss rates (respectively) in a sir1Δ background for loci with different nucleosome number (different genetic length). The results from the model are shown in black and, in red, the experimental data from Ref. (Saxton & Rine, 2019). C,D) Switching rates shown as in A and B, but shown for a wider range of genetic lengths, by combining data from Ref. (Saxton & Rine, 2019), in red, with new experimental data obtained specifically for this study (blue). The labelling for panel D is the same as for panel C. There are two datapoints for the 14 nucleosome strains, as two different strains were created by introducing the same DNA into two different locations at the locus. The results of the model are shown in black, but the parameters have been modified slightly with respect to the results in A and B, to better capture the new data. For details regarding the parameters, see SI Appendix, Section 1.6. The error bars represent the 95% confidence interval.

There are two effects at play when reducing the number of nucleosomes between the silencers in the simulations. First, since we still have two silencers, their effect on the remaining nucleosomes is stronger (the number of nucleosomes per silencer decreases), making the locus more prone to silencing. Second, as pointed out above, there is a decrease in the reliability of inheritance for smaller loci. In the case of the silencing loss rate, these effects counteracted each other to a certain extent: as the locus size was reduced, the silencers became more effective (lowering the silencing loss rate), but inheritance less faithful (increasing the loss rate). Crucially, for the case of silencing establishment, both effects reinforced this rate as we decreased the nucleosome number, rather than cancelling out, leading to qualitatively different behaviour.

The model predicted a continuation of this trend for larger loci, an important result which we therefore tested experimentally, since any such trend identified in earlier experiments was not statistically significant. Strains with 14 and 16 nucleosomes in the *HMR* locus were created by inserting intergenic DNA found at *HML* into the endogenous *HMR-GFP* locus. We tested other sources of intragenic DNA with similar results regarding the overall fraction of silenced cells (see *SI Appendix, Supplementary Figure 1*), since every mutant with an increased number of nucleosomes at the locus showed a higher degree of fluorescent protein expression than for the wild-type length (12 nucleosomes), as expected from the model.

To test the model predictions, we experimentally measured the switching rates for these strains and, in agreement with the model, the silencing establishment rate decreased substantially (Fig. 2C), a decrease that was statistically significant (see *SI Appendix, Supplementary Figure 2*). In fact, the decrease in the silencing establishment rate was even more marked than initially predicted by the model, which forced us to modify certain parameters. Since the effect of silencers is partially responsible for the trend in the silencing establishment, giving more importance to silencers in the model (with respect to nucleosomes) should increase the steepness of the trend. By increasing the parameter that controls the affinity of the SIR complex to silencers and decreasing that for histones, the model was able to better capture the 20-fold drop in the experimental establishment rate over the 6 to 16 nucleosome range (Fig. 2C, for details on the parameters used in Fig. 2C and D and the rest of the paper see *SI Appendix, Section 1*.*6*, and for the results with the original parameter set see *SI Appendix, Supplementary Figure 3*).

With respect to the silencing loss rate, the model predicted limited changes with both parameter sets (Fig. 2B and D). The experiments confirmed this as the silencing loss rate did not change significantly when increasing the genetic length of the locus (Fig. 2D and see *SI Appendix, Supplementary Figure 2*). Overall, there did not seem to be a trend in the silencing loss rate since there were no statistically significant changes within the two experimental datasets. Nevertheless, there was a substantial dispersion between the results obtained in this work and those of Ref. (Saxton & Rine, 2019) for reasons that were unclear, although this ∼2-fold dispersion remains very small compared to the 20-fold variation in the establishment rate. Altogether, the model predictions for silencing establishment/loss rates as a function of locus size were well supported by our new experimental data.

### Chromatin binding

As argued above, silencing at the *HMR* locus occurs in a highly cooperative manner, due to the SIR complex binding and H4K16ac feedback. Traditionally the process had been viewed as nucleation of the Sir proteins starting at the silencers and subsequent spreading towards the rest of the locus. In contrast, in Ref. (Saxton & Rine, 2022), two of us found that the extent of Sir binding on nucleosomes also has an effect on the occupancy levels of the silencers, making cooperativity between silencers and nucleosomes a two-way interaction.

The model has no sequence dependency other than the silencers’ position and considers every nucleosome to be able to equally interact with any other. Thus, the extent to which such a model could capture the observed cooperativity between genetic features in the locus was unclear. By reanalysing ChIP data for SIR complex binding in different histone mutants, an overall nucleosome occupancy value was obtained, which could be compared to the outcome of the model. Fig. 3A depicts the Area Under the Curve (AUC) for the E silencer as a function of the AUC of the nucleosomes across the locus for the different histone mutants, with both quantities normalised to the wild-type. This analysis revealed a good accordance between the experimental data and the model. The histone and Sir mutants were chosen to reduce the spreading on nucleosomes but not directly alter the binding to silencers. These included *H4K16Q* (an acetyl mimic), *H3K79L* (a methyl mimic), *Sir3-bahΔ* (Sir complex whose nucleosome-binding domain has been deleted) and *sir2-N245* (catalytically inactive for H4K16ac deacetylation). For details on the modelling of these mutant strains, see *SI Appendix, Section 4*. In agreement with the mutual cooperativity picture between silencers and nucleosomes, as the Sir binding to nucleosomes decreased, so did the E silencer binding, both in the model and the experiments. Note the non-linear cooperativity, with more than half of the drop in E silencer binding taking place when the nucleosome Sir binding is less than 20% of the wild-type value.

**Figure 3.**
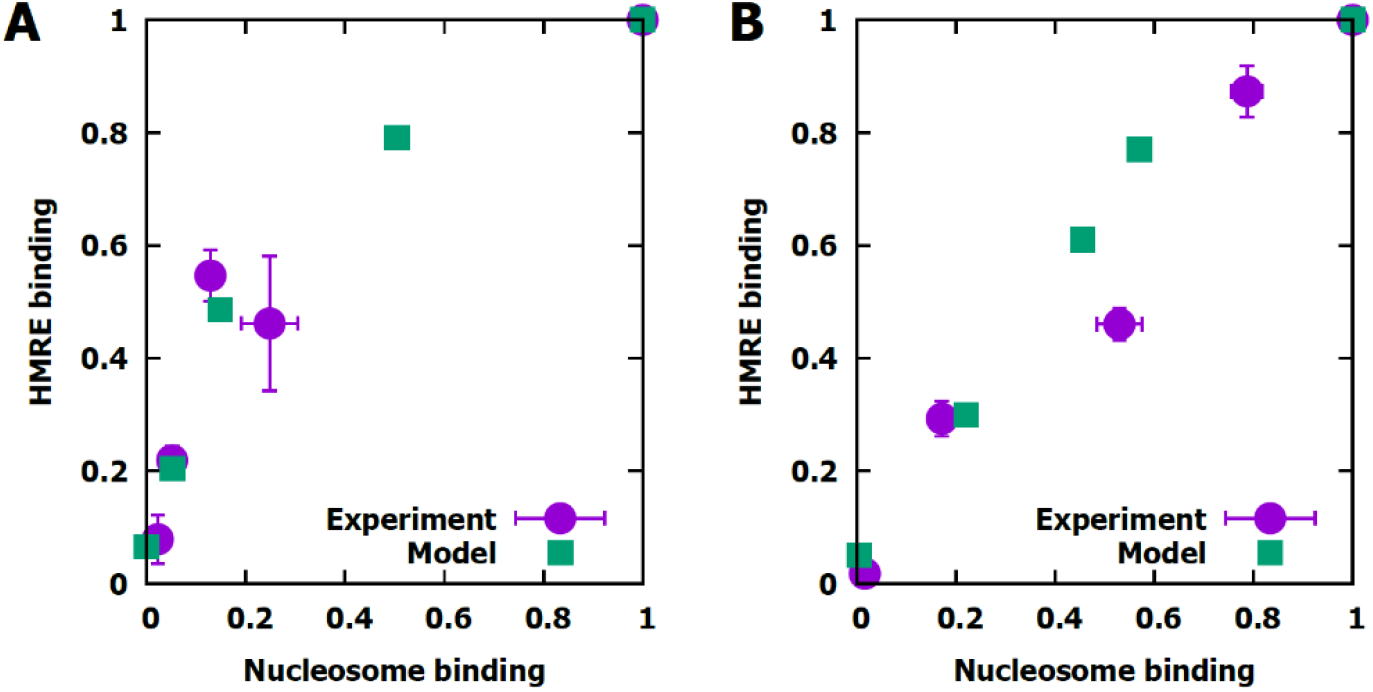
Binding of the SIR complex to the E silencer and to nucleosomes. A) Binding at the E silencer (HMRE) as a function of binding at nucleosomes for different histone mutants. Experimental data from (Saxton & Rine, 2022) is shown in purple circles and modelling, in green squares. The data (modelling and experimental) is normalised to the wild-type, shown in the top-right hand corner. The mutant strains are, from right to left, H3K79L, H4K16Q, sir3-bahΔ and sir2-N345. B) Same data as A but for a Sir4 titration. The experimental data (strain with estradiol-inducible promoter) is normalised to the population with the highest SIR complex binding in the titration (data from Ref. (Saxton & Rine, 2022)) while the modelling output has been normalised to the wild-type Sir concentration in simulations. Experimental error bars correspond to standard deviation (independent in each direction) from two different replicates.

According to the model, the cooperativity signature seen in these experiments resulted from the way the SIR complex binds to the locus (dimerisation) and the deacetylation activity of Sir2. As the amount of Sir binding at the nucleosomes decreases, so does the number of unmodified H3-H4 pairs (due to lower deacetylation rates). This results in a lower binding of the SIR complex to the E silencer, as it requires two binding sites, one of them being the E silencer and the other one being either the I silencer or the depleted unmodified H3-H4 pair pool. This argument highlights the importance of the silencer-nucleosome cooperativity and explains why most of the drop in E silencer binding occurs when nucleosome Sir binding is low: the E silencer only needs one unmodified H3-H4 pair to dimerise with, hence until the number of unmodified H3-H4 pairs becomes very low the E silencer does not feel much of the effect.

In a similar manner, we show the E silencer binding against nucleosome binding in a Sir4 titration, using an estradiol-inducible promoter to tune the concentration (data from Ref. (Saxton & Rine, 2022), see Fig. 3B). This time the relationship was roughly linear in both simulations and experiments, since we were varying the overall SIR complex concentration in solution, which directly affected silencers and nucleosomes equally.

Taken together, the analysis shown in Fig. 3 implied that the minimal mathematical model used for chromatin, which represents short-lived locus-wide contacts, was adequate and could capture the observed cooperativity between silencers and nucleosomes. In addition, the model shed light onto the origin of the silencer-nucleosome cooperativity, establishing as a main cause the availability of unmodified H3-H4 pairs with which the silencer can dimerise.

### Switching rates in Sir4 titrations

In addition to the ChIP profiles used in Fig. 3B, switching rates were obtained at different estradiol concentrations (Saxton & Rine, 2022), thus modulating the Sir4 concentration and the switching properties of the locus. To compare the results from experiments with that of the model, we measured the switching rates in the simulation when varying the Sir complex concentration. However, the titrated Sir4 and native promoter Sir4 switching rates were different even at approximately the wild-type concentration, possibly due to noise introduced by the inducible promoter. To keep the modelling within a common framework, we therefore multiplied the theoretical switching rates by the ratio between the switching rates from the estradiol-inducible promoter and the native promoter. In addition, a linear conversion was made to estimate the SIR complex concentration in the simulations from the estradiol concentration in experiments (for details see *SI Appendix, Section 5*). After this linear transformation, there was good agreement between model and experiments (see Fig. 4A).

**Figure 4.**
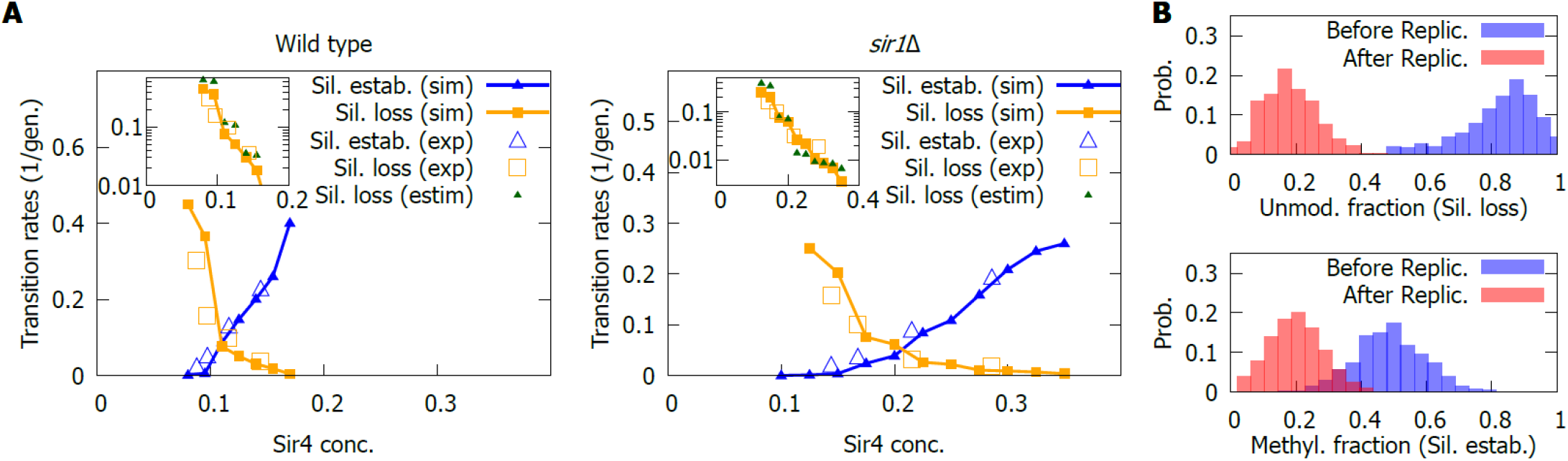
Switching rates in a Sir4 titration. A) Silencing establishment and silencing loss rates in in the wild-type (left) and in a sir1Δ background (right). Insets show the silencing loss rates, in logarithmic scale, including an estimate (estim.) based on the inheritance probabilities at chromosome replication, see Eq. [2]. The experimental data was obtained from Ref. (Saxton & Rine, 2022) and was normalised to match switching rates from the native promoter, to keep modelling within a common framework. B) Histone PTMs before (blue) and after (red) replication. Top: Fraction of unmodified H3-H4 pairs before and after the replication event at the beginning of the generation where silencing loss occurs. Bottom: Fraction of methylated H3-H4 pairs before and after the replication event at the beginning of the generation where silencing establishment takes place. Results in panel B shown for a sir1Δ mutant at wild-type concentration.

As expected, both in the wild-type *SIR+* and *sir1*Δ mutant, experiments and modelling agreed on a decrease in the silencing loss rate and increase of the establishment rate as the Sir4 concentration rises. Importantly, the region of bistability of the wild-type silencers is at low Sir concentration, while for the *sir1*Δ strain, it is at higher concentrations.

More unexpected was the way in which the switching rates vary with Sir concentration. The model predicted a roughly exponential trend for the decrease of the loss rate (see insets in Fig. 4A), consistent with the experimental data. However, this is not the case for the establishment rate (see insets in *SI Appendix, Supplementary Figure 4*), both in the model and the experimental data, hinting at an asymmetry in the way the switching occurs.

One intuitive reason behind this asymmetry could be the effect of replication. In the simulations, when a cell was silenced at *HMR*, most H3-H4 pairs at *HMR* were unmodified and many of them were substituted by acetylated pairs at replication, while in a cell lacking silencing, acetylated and methylated histones were substituted by only acetylated ones. Thus, to obtain insights into the contribution of replication dynamics on the switching of transcriptional states, the distribution of unmodified histones before and after replication was obtained (shown in Fig. 4B, top), only for cases where *HMR* in the daughter cell became expressed after it was not in the mother cell, i.e., silencing was lost. On average, only half the unmodified H3-H4 pairs should have been lost. However, in cases where the silencing was lost after replication, substantially more than half were lost in most cases, implying a critical role for replication in the loss of *HMR* silencing.

Therefore, one could speculate that most of the stochastic contribution to the silencing loss stems from an atypically large loss of unmodified nucleosomes at replication and not from the dynamics of the system after replication. In that case, one could set a threshold for the maximum number of unmodified H3-H4 pairs a locus can have in order to lose silencing in the following generation. If, after replication, the locus possessed more unmodified histone pairs than the threshold it would remain silenced, but it would otherwise become expressed. The following approximation to the silencing loss rate could then be obtained, given the knowledge of the distribution of unmodified H3-H4 pairs in the silenced state before replication *P*_sil_(*n*_*u*_) and the threshold *M*:

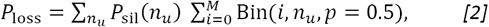

Where *P*_loss_ is the silencing loss probability per cell and generation and Bin(*i, n*_*u*_, *p* = 0.5) is the binomial probability distribution of *i* unmodified H3-H4 pairs remaining after replication, given that there were *n*_*u*_ before, and that the probability *p* of losing each of them is 50%. The sum over *i* implies that, in this approximation, any locus with a number of unmodified H3-H4 pairs between 0 and *M* will become expressed. The sum over *n*_*u*_ accounts for the probabilities that the locus has a different number *n*_*u*_ of unmodified H3-H4 pairs and, thus, it would require a different number of successful removals of unmodified histones to become activated. The threshold *T* was obtained from the average of the probability distribution after replication for cells that will become expressed (red probability distribution in Fig. 4B, top).

This estimate for the silencing loss probability involved several approximations and assumptions: that the stochastic contribution stems exclusively from replication and not from any subsequent histone PTM dynamics and that histone pair removal at replication is independent of each other (a condition that explains the appearance of the binomial distribution in the estimate). As can be seen from the inset in Fig. 4A, these assumptions led to an estimate that was often off by a ∼50% with respect to the full model, but did set correctly the order of magnitude of the silencing loss rate and recapitulated the exponential trend. Therefore, the value of this estimate is more qualitative than quantitative: it suggested that the main contribution to the silencing loss rate was a rare replication event, where more unmodified histones than average fail to be inherited, but where other factors also have an effect in the silencing loss rate, which explains why the quantitative values are not accurate. Intuitively, the exponential trend was due to replication-dependent nucleosome removal being an independent event for each nucleosome, which is reflected in an exponential factor from the binomial distribution if the threshold scales linearly with the Sir4 concentration. In fact, the threshold did not decrease in an exactly linear manner with the concentration, but this deviation was partially corrected by the combinatorial prefactor in the binomial distribution.

When this approach was repeated with the silencing establishment rate, the resulting estimate failed to reproduce the outcomes of the model (see *SI Appendix, Supplementary Fig. 4*). This result implied that, for the silencing establishment, the dynamics of histone PTMs during the rest of the cell cycle are a major source of stochasticity. This could also be seen from the distribution of methylated histones before and after replication (as methylation is the sole activating mark that can be lost at replication), in cases where silencing will be established in the next generation. In Fig. 4B (bottom), these distributions are shown, and the average methylation loss is roughly half of the total amount (the expected average), suggesting that silencing establishment was not very dependent on loss of methylation at replication.

Overall, the model was able to recapitulate the observed switching rates at different Sir concentrations and identified two different potential mechanisms for the transitions: a mechanism based on an anomalously large loss of unmodified histones at replication for the silencing loss, and a stochastic fluctuation in the histone PTM kinetics for the silencing establishment. In addition, the model predicted an exponential trend for the silencing loss that was consistent with the experimental data.

### Bistability of silencing in different mutation backgrounds

Another useful aspect of the Sir protein titrations is the ability to look for bistable behaviour for concentrations that would otherwise not be experimentally accessible. Some Sir4 titrations for mutants affecting silencer strength were reported in Ref. (Saxton & Rine, 2022), leading to the conclusion that mutants in which silencers are made weaker are bistable at certain Sir4 concentrations, but even strong-silencer strains, such as the wild-type, become bistable with low enough Sir4 concentrations. Silencer strength was modulated by removing proteins that bind exclusively to silencers (such as Sir1) or by removing binding sites for scaffolding proteins at the silencers (such as the Rap1 binding site, Rap1 b.s.).

The model recapitulated the fact that bistability is enhanced in a different range of Sir concentrations in each strain, depending on the strength of the silencers. By lowering the concentration of Sir proteins, we obtained loci that would transition from a fully expressed state (mostly acetylated) when the Sir concentration is low, to a fully silenced one for high Sir concentrations, passing through an intermediate concentration region where the system can transition back and forth between active and silenced states (see *SI Appendix, Supplementary Figure 5*). We quantified the bistability of the model for each silencer strength mutant at various concentrations using the bimodality coefficient (BC), defined in Ref. (Institute, 2012). The BC is a function of the skewness and the kurtosis of the distribution (of acetylated histones, in this case), is positive and bounded from above by 1 (for details on the implementation, see *SI Appendix, Section 6*). Typically, values of the BC larger than ∼0.55 are considered to correspond to bimodal distributions, which is an indicator of two stable states. The model predicted a peak in the BC for all mutants considered, but the location of the peak shifted towards smaller Sir4 concentrations for stronger silencers, see Fig. 5, red triangles.

**Figure 5.**
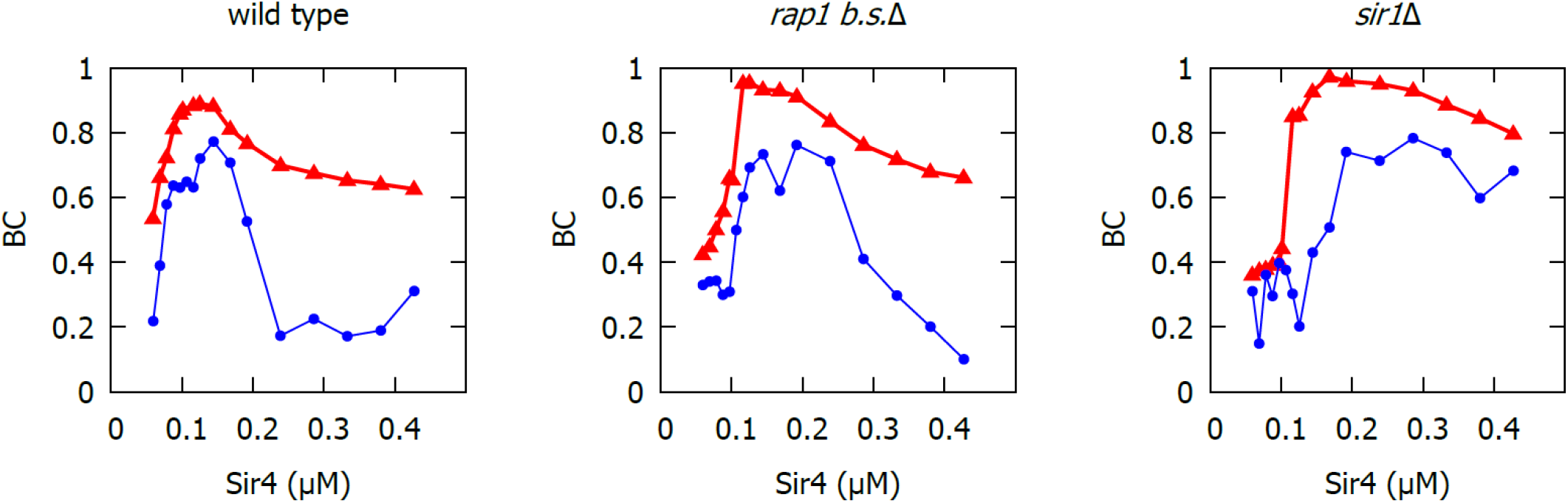
Bistability in a Sir4 titration. Bimodality coefficient (BC) for the wild-type and two silencer strength mutants, at different Sir4 concentrations. Experimental BC in blue (circles), theoretical BC in red (triangles). Experimental data from Ref. (Saxton & Rine, 2022).

We then sought to compare the theoretical predictions with the experimental data, by computing the BC from the distribution of GFP intensity (in logarithmic scale, as reported in (Saxton & Rine, 2022)) obtained from experiments. The comparison between the experimental and the theoretical BCs, for different silencer strength mutants and Sir4 concentrations, is shown in Fig. 5.

While the comparison was not quantitatively precise, the overall trends and location of the peaks were captured by the model, showing that stronger silencers tend to have a peak in bistability at very low Sir4 concentrations, but weaker silencers can broaden the bistability window and shift it towards higher Sir complex concentrations. It is remarkable that the *sir1Δ* strain could sustain a high bistability degree over a large concentration window, both in experiments and in the model, as opposed to *rap1 b*.*s*.*Δ* or the wild-type that displayed a more pronounced peak at mid or low concentrations.

Reasons for the imperfect quantitative agreement may include using the same methodology (except for the logarithmic scale in the fluorescence assays) for experimental and synthetic data (which have different sources of noise) or the metric being heavily impacted by very rare events which were either not seen in experiments or overestimated by the model. Finally, given that the output of the model was at the level of chromatin modifications, but the output of experiments was fluorescent protein levels, another reason for a reduced quantitative agreement could be that transient fluctuations at the chromatin level need not translate to fluctuations at the protein level, yielding a lower experimental BC.

## Discussion

In the present work we have developed the hypothesis of dynamic changes in the three-dimensional conformation of the locus substantially affecting the removal of histone modifications, thus bidirectionally linking chromatin compaction and histone PTMs. Deacetylation of H4K16 compacts the locus, which promotes Sir binding and further deacetylation via read-write activity of Sir2 with H4K16ac. Hence, we favoured a picture whereby compaction is both cause and consequence of the histone PTM state of the locus: it is consequence as the chromatin conformation resulting from a silenced, unacetylated locus is compact, and it is cause since compact chromatin increases the rates of enzymes, such as the Sir2 deacetylase, that require contacts within the locus for their activity.

Not only is this an attractive hypothesis, but it is also solidly grounded on the current biochemical data. Recent studies found that cooperativity of Sir proteins is limited to a dimerisation process (Behrouzi et al., 2016), suggesting that Sir binding alone cannot explain the highly cooperative behaviour observed in the dynamics of HMR. Indeed, previous modelling studies had captured the bistable behaviour of the *HML* locus (Sneppen & Dodd, 2015), but only by requiring much more cooperative protein binding dynamics than is observed. Thus, we identified another source of cooperativity, linking the acetylation of H4K16 and the activity of Sir2 to considerable changes in chromatin compaction, which have been observed *in vitro* (Johnson et al., 2009). Despite its relatively simple form, the model captured and helped elucidate the nature of the cooperativity in the *HMR* locus, which is a hallmark of epigenetic silencing. From this, we concluded that the experimental data were consistent with a mechanism of locus-wide cooperativity that is dominated by the dimerization behaviour of the Sir complex, often between unmodified nucleosomes and silencers far away in genetic distance, but close in the physical distance. Long distance interactions such as these ones had been reported before (Valenzuela et al., 2008), but the extent to which they contributed to the overall cooperativity in the locus had remained unclear.

Another key finding was the asymmetry in the behaviour of silencing loss and establishment rates when subjected to changes in Sir4 concentration. According to the model, these two events had different origins; the silencing loss probability was mostly controlled by the stochasticity of nucleosome replacement at replication, while for silencing establishment the stochastic histone PTM dynamics played a major role.

Moreover, the switching rates at a variety of locus lengths were well captured and new behaviour predicted by the model was experimentally verified, highlighting the importance of size and physical space in genetic regulation. Two metrics were especially relevant: the one-dimensional genetic length between the silencers and the three-dimensional conformation of chromatin. If the genetic length was reduced, then the silencers would have a greater influence on the remaining (fewer) nucleosomes. In addition, a compact locus in three-dimensional space would enhance Sir binding and Sir2 deacetylation (which depend on intra-locus contacts), but a locus in an expanded conformation would allow other PTMs to take over (H4K16ac and H3K79me3, in this case). Thus, the model successfully integrated one and three-dimensional effects into the dynamics of the locus, capturing crucial features of the locus and making predictions that were experimentally verified. This goes beyond previous theoretical work focused on explaining the interplay between epigenetic marks and chromatin folding (Michieletto et al., 2016; Nickels et al., 2021; Owen et al., 2022; Sandholtz et al., 2020) by providing detailed mechanistic insights and quantitative predictions that can be experimentally tested.

Overall, we have found that the dynamic compaction and decompaction of the locus underpins the bistability of the locus in silencer strength mutants and maintains a robust and heritable transcriptional state across tens to hundreds of generations. However, it remains to be explored how robust it is with respect to cell-cycle dependent processes other than DNA replication, such as genome-wide acetylation changes (Wilkins et al., 2014). Another limitation of this work was the isolated description of the HMR locus in the model: placing the locus in the context of the chromosome and comparing the regulation of the locus with that of other (possibly active) genes would yield valuable insights (Nickels & Sneppen, 2023). Finally, if more detailed measurements of the conformation of the locus were available, a more ambitious physical modelling could be attempted (Farr et al., 2021), which would most likely improve the predictive power of the model.

Taken together, in this study we have proposed and developed a role for chromatin compaction, in which compaction is tightly associated with histone modifications, influencing each other as part of a unified regulatory mechanism. This broader view complements the classical view of chromatin compaction as downstream of histone PTMs and regulatory circuits, by merely blocking transcription sterically. Our model and data, as well as previous literature, allow us to paint a more complete picture of the epigenetic regulation of the HMR locus in budding yeast, underscoring the importance of chromatin conformation and the physical dimension of the problem. We expect that the concepts developed here, such as the two-way feedback interaction between acetylation and chromatin compaction, may be relevant in many other gene regulatory circuits.

## Materials and Methods

### Model and implementation

A full description of the model, including all the details and parameter values can be found in the *SI Appendix, Sections 1, 2 and 4*.

The implementation of the model, as well as the data generated for the figures in this paper, can be found in https://github.com/AMovillaMiangolarra/Conformation_Acetylation_Feedback_2023. The statistics shown in the figures of this work correspond to simulations sampled over 10^5^ generations.

### Experimental methods

#### Strain construction

Strains and oligonucleotides used in this study are listed in Table S1 in the *SI Appendix*. Insertions of DNA into *HMRα* were achieved using CRISPR/Cas9 technology (Lee et al., 2015). To do this, sgRNAs that spanned the precise insertion sites were generated. Additionally, repair templates containing DNA from *HML, K*.*lac Leu2*, or *K*.*lac Ura3* were amplified with flanking homology to *HMRα*, such that recombination of these templates with *HMRα* would insert DNA within the sgRNA cut site. Co-transformation of a Cas9/sgRNA plasmid with an appropriate repair template produced cells that constitutively cut the sgRNA-targeted site and thus remain arrested; these cells would only grow into a colony if a mutation, such as insertion of the desired repair template, disrupted the sgRNA-targeted site. All mutations were confirmed by sequencing.

#### Flow cytometry

To allow bistable populations of cells to reach equilibrium prior to flow cytometry, cells were grown at log-phase for 24 hours. More specifically, three separate colonies (representing three technical replicates) of a given strain were used to inoculate three different cultures in YPD liquid media, which were grown to saturation at 30°C overnight. These cultures were then serially back-diluted in YPD liquid media and maintained at log-phase at 30°C over 24 hours. After 24 hours of log-phase growth, cells were pelleted, resuspended in 100 μL of 4% paraformaldehyde, fixed for 15 minutes at room temperature, pelleted, and then resuspended in 150 μL of 1x PBS solution.

Flow cytometry was performed with an LSR Fortessa (BD Biosciences) with a FITC filter for Green Fluorescent Protein (GFP). At least 30,000 cells were analyzed per replicate. FlowJo software (BD Biosciences) was utilized to analyze flow cytometry data. All cells in the experiment were gated identically.

#### Microscopy

Prior to microscopy, cells were grown in YPD overnight at 30°C, and then to log-phase in the same conditions. 3 μL of 0.5 OD cells were then spotted onto CSM agar plates. Once the spots were dry, a sterile spatula was used to cut out a small square that encompassed the cell spots, which was then removed and inverted onto a 35 mm glass bottom dish (Thermo Scientific 150682). Cells were then imaged using a Zeiss ZA inverted fluorescence microscope with a Prime 95B sCMOS camera (Teledyne Photometrics), Plan-Apochromat 63x/1.40 objective (Zeiss), MS-2000 XYZ automated stage (Applied Scientific Instrumentation), and MicroManager imaging software (Open Imaging).

To generate timelapse images, samples were kept at 30°C and humidified by a P-set 2000 Heading Incubation Insert (PeCon). Brightfield and fluorescence images were collected for 16 fields of view per sample, every 20 mins for 10 hrs. A single Z-slice was acquired for each field of view at each timepoint. Images were analyzed with ImageJ (NIH). Silencing loss and silencing establishment events were counted manually with a single-blind approach.

## Acknowledgements

We thank all members of the Howard group for discussions. We also acknowledge financial support from BBSRC Institute Strategic Programme GEN (BB/P013511/1) to MH and a grant from the National Institutes of Health (R35GM139488) to JR.

## Supplementary Information Appendix for

### 1 Model

The model describes the state of a varying number of nucleosomes (between 6 and 16). Nucleosomes contain two copies of the histone proteins H2A, H2B, H3 and H4; forming two H2A-H2B heterodimers and a (H3-H4)_2_ heterotetramer, but only the PTMs in the (H3-H4)_2_ heterotetramer seem to be relevant for HMR silencing [1]. The DNA wraps around the nucleosome, exposing two faces of it, each of them containing a single copy of the histone proteins. While allosteric effects can influence the deposition of histone marks within a single face of the nucleosome (e.g. [2]), to our knowledge there is little or no evidence of allostery across the entire nucleosome. Then, each face of the nucleosome is independent of the other [3], allowing PTMs (or Sir proteins) to establish independently in each of the copies of the histone proteins that constitute the same nucleosome. Thus, the model describes the state of pairs of H3-H4 histones independently (representing one face of the nucleosome). These H3-H4 pairs can be found in five different states:

- **Unmodified**: They allow for SIR complex binding and dimerization and thus, can be sub-divided into two different situations:
  - Bare (U): With no Sir bound
  - Bound (Ub): The SIR complex is bound to the unmodified H3-H4 pair. This could be by dimerising with another H3-H4 pair or with a silencer. We do not consider the case of isolated SIR-bound dimers since Sir3 seems to have a much lower affinity for them [3].
- **Acetylated** (A): H4K16 acetylation. There is no H3K79 methylation in this state.
- **Methylated** (M): H3K79 methylation and no H4K16 acetylation.
- **Acetylated and methylated** (AM): H3K79 methylation and H4K16 acetylation.

We assume that all nucleosomes interact with every other nucleosome in the locus as a consequence of a rather compact and dynamic three-dimensional arrangement. In our simulations, this implies that every H3-H4 pair will be able to interact with every H3-H4 pair. Therefore, we do not need to keep track of the spatial positions of each nucleosome or H3-H4 pair (spatial mean-field approximation) and *L* is the number of H3-H4 pairs in the locus (twice the number of nucleosomes), which satisfies *n*_*m*_ + *n*_*a*_ + *n*_*am*_ + *n*_*u*_ + *n*_*ub*_ = *L*, where *n*_*x*_ is the number of H3-H4 pairs that are methylated (*x* = *m*), acetylated (*x* = *a*), both methylated and acetylated (*x* = *am*), unmodified (*x* = *u*) and Sir-bound (*x* = *ub*). Note that this approximation, where all nucleosomes interact with all others, might become inaccurate for larger loci, where this is no longer physically feasible. Based on Ref. [3], the dynamics of Sir3 proteins seems to be on the timescale of tens of seconds. According to Ref. [4], methylation and epigenetic states seem to survive for at least a few generations (timescale of hours). Then, we can classify the processes in the system into two different timescales: a fast one for Sir protein binding and a slow one for histone modifications. Consequently, in the model, the rates of the processes in the fast timescale (Sir protein binding, unbinding or dimerisation) will include a factor *τ*_*s*_^−1^, where *τ*_*s*_ ≃ 10s, marking the processes as fast when compared to the histone PTMs.

If the the H3-H4 pair is unmodified, Sir proteins will be able to bind to the surface of the nucleosome. This will not be the case if the H3-H4 pair is either acetylated, methylated, or both. However, even if an H3-H4 pair is acetylated or methylated, they will still be able to bind the opposite surface of the nucleosome, as long as its corresponding H3-H4 pair is unmodified. For simplicity, we model the binding of Sir proteins (the SIR complex, Sir2-4) to nucleosomes as a single step.

Silencers are modelled as H3-H4 pairs which are always in the unmodified state, thus always able to dimerize and recruit SIR proteins, but they have an affinity for Sir proteins larger than that of nucleosomes. In the *sir1 Δ* mutant, for simplicity and roughly in agreement with ChIP experiments [5], we take the affinity of the Sir complex to silencers to be almost as low as for an unmodified nucleosome.

To mathematically model this system, we still need to determine the reaction rates. This will be done in the following sections.

#### 1.1 Fast timescale: Sir binding to chromatin

In Ref. [3] the cooperativity between Sir proteins binding to nucleosomes is analysed, which until then had been a source of controversy [6]. By means of electrophoretic mobility assays, they were able to infer that indeed there is a cooperativity in the binding of Sir proteins (Sir 3 and 4) to nucleosomes but that it is limited to one interaction with a single other protein. This is, the process of Sir binding to chromatin resembles that of dimerization and occurs on a timescale of tens of seconds, which is much faster than the cell cycle (few hours). Importantly, they found that, *in vitro*, the dissociation constant between the nucleosome and a single Sir3-4CC (Coiled Coil) protein is *K*_D_ ≃ 1.4*μ*M, which is a higher concentration than the one estimated for Sir3 in vivo (≃ 0.5*μ*M [3, 6]). However, the cooperative binding of two Sir proteins to two different nucleosomes had a much lower dissociation constant (*K*_D_ ≃ 0.08*μ*M), implying that the nucleosome dimerization is the primary binding pathway. Therefore, in the model, we consider that the binding of Sir proteins to nucleosomes occurs only in pairs (or dimers) and that they only bind unmodified nucleosomes, neglecting any affinity for acetylated or methylated nucleosomes.

In addition, the HMR locus is flanked by two silencers (protein binding regions) with a strong affinity for Sir1 and the rest of the SIR complex. We thus assume that the SIR complex can also bind to the silencers, dimerising in a similar way with the rest of the locus (the affinities for these dimerisation reactions will be discussed below). Hence, the Sir binding to chromatin is described by the vector (*n*_*ei*_, *n*_*en*_, *n*_*in*_, *n*_*u*_, *n*_*d*_), where *n*_*ei*_ = 1 if the silencers are bound between themselves (0 otherwise). If the SIR complex is bound to the E or I silencer and to an unmodified nucleosome then *n*_*en*_ = 1 or *n*_*in*_ = 1, respectively (0 otherwise). Finally, *n*_*d*_ corresponds to the number of H3-H4 histone pairs dimerised by the SIR complex, while *n*_*u*_ is the number of unmodified H3-H4 histone pairs to which Sir proteins are not bound. Thus, the vector

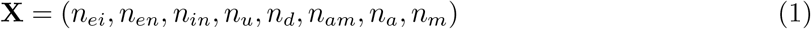

describes the full state of the system including the silencers, protein binding and the histone post-translational modifications.

There are four relevant reactions describing the dynamics of Sir binding to the locus:

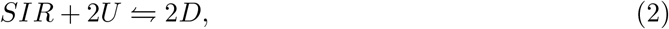

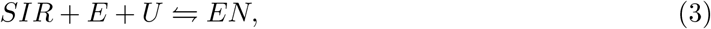

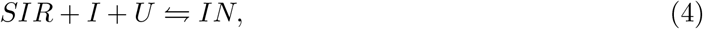

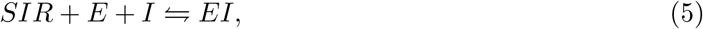

whose dynamics, in the all-interacting locus approximation, follow mass action kinetics [7]. Then, in continuous time modelling (see Section 2 for mathematical details of the simulation) the forward rates (left to right direction) take the following form:

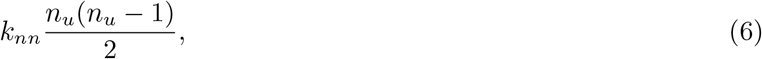

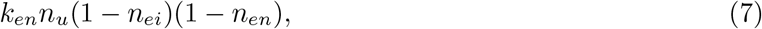

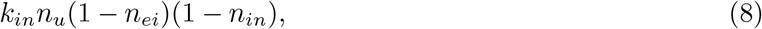

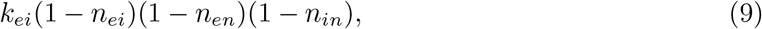

where factors of the form (1 − *n*_*ei*_)(1 − *n*_*xn*_) ensure that there is only one Sir complex that can bind to the *X* silencer, where *X* is either *E* or *I*. The backward rates are taken to be

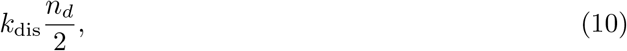

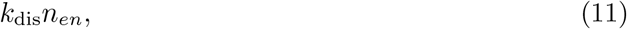

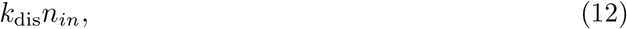

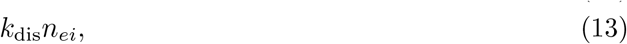

where, for the sake of simplicity, it has been assumed that the unbinding rates are all equal. Given that these processes occur within the fast timescale of protein binding to chromatin, an appropriate choice of the backward reaction constant is *k*_dis_ = *τ* _*s*_^−1^.

In order to choose the forward reaction constants in a coherent way, the reaction constants were constructed by assuming that they are a product of the affinity of the two anchoring points of the SIR complex:

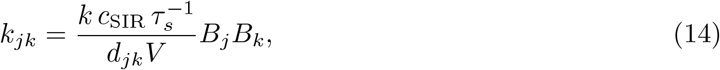

where *c*_SIR_ is the concentration of the SIR complex in solution in the nucleoplasm (which we take to be a constant) and *jk* = *ei, en, in, nn* depending on whether the SIR complex binds to the E silencer, the I silencer or a nucleosome, respectively. Consequently, the magnitude of the parameter *B*_*i*_ represents the strength of the bond formed between the chromatin anchoring point and the SIR complex. The parameter *k* is the characteristic constant of the binding by dimerisation reaction and is divided by an approximation of the typical three-dimensional volume in which the corresponding particles are diffusing, *d*_*jk*_*V*.

The three-dimensional volume is computed by assuming the Flory picture used to estimate the overall volume of the locus (see section 3) is still valid for distances within the locus. This theoretical estimate yields that the volume is proportional to the genetic length, but different particles will be at different genetic lengths from each other. For example, the silencers will be at a distance *∝ L* between themselves, while the silencers are, on average, at a distance *∝ L/*2 from nucleosomes and nucleosomes are at an average distance *∝ L/*3 between themselves (see below). Thus, the denominator in eq. (14) is taken to be the product of the overall locus *V* and a factor *d*_*jk*_, which takes into account these differences in genetic distance between silencers and nucleosomes. They take the following values:

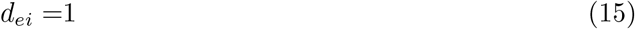

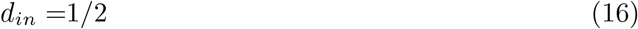

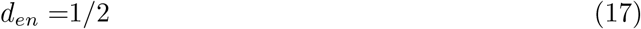

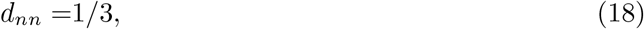

in accordance to the argument given above. Then, the binding constant for silencer-nucleosome dimerisation is 2*/*3 of that for nucleosome-nucleosome, and the silencer-silencer constant 1*/*3 of that for nucleosome-nucleosome, as stated in the main text.

The values of the rest of the parameters are specified in Section 1.6.

##### 1.1.1 Calculation for the average nucleosome-nucleosome genetic distance

Given *L*_*n*_ = *L/*2 nucleosomes stacked in a straight line, the average nucleosome-nucleosome distance (assuming unit distance 1 between each consecutive one) is:

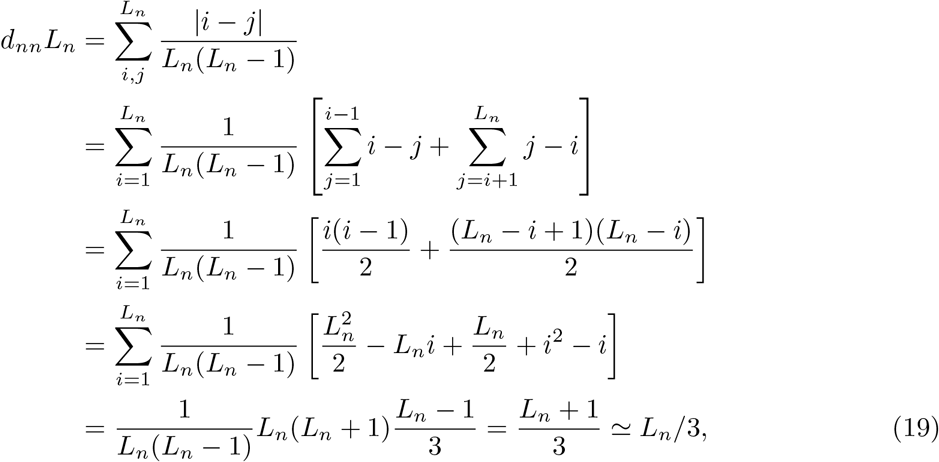

where the first three terms of the second to last line have cancelled each other, and the last approximation is valid if *L*_*n*_ *≫* 1.

#### 1.2 Sir proteins concentration and titrations

We have so far described the binding of the Sir complex as a whole to chromatin. However, the Sir complex is formed by three proteins (Sir2-4). Sir3 is key for the complex to bind nucleosomes and its concentration has been estimated to be roughly 0.5*μ*M [3, 6]. Given that a Sir complex requires a Sir3 dimer we assume the baseline concentration of the complex *c*_SIR_ ∼ 0.25*μ*M.

However, in the experiments of Ref. [5], the concentration of Sir4 is varied by means of an inducible promoter that can be tuned varying the concentration of estradiol. To mimic this case, in the model we vary linearly the concentration of the entire SIR complex in solution:

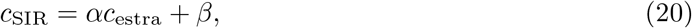

where *c*_estra_ is the concentration of estradiol, *α* is a constant relating the SIR complex concentration to the estradiol concentration (via Sir4 modulation) and *β* is a small (positive) correction, which could mimic a low basal noisy transcription in the absence of estradiol.

As a first approximation, this is a valid approach since, if Sir4 is limiting for the creation of the complex and there is Sir3 in excess, one would expect the overall concentration of the complex to increase linearly with the concentration of Sir4. Nevertheless, as we increase the concentration of Sir4, it will stop being limiting and this linear approximation may fail.

#### 1.3 Post-translational modifications

On top of the binding dynamics described in Section 1.1, the simulation includes stochastic histone post-translational modification dynamics and replication. We consider three types of enzymatic reaction that can occur at the locus:

- **Sas2 acetylation**. Sas2 is the catalytic subunit of the yeast histone acetyltransferase SAS (Something About Silencing) complex, which preferentially acetylates H4K16. Sas2 is thought to have more affinity for histones in solution, but it can also acetylate histones incorporated in chromatin [8]. We model the acetylation reaction at the locus with a propensity function linear with the amount of substrate available for Sas2 (that is unacetylated H3-H4 dimers):

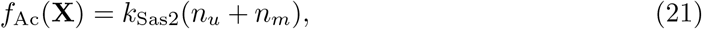

where *k*_Sas2_ is the characteristic rate constant of Sas2 acetylation of histones. Here we have two terms corresponding to the two types of histones Sas2 can acetylate: either unmodified H3-H4 dimers (without Sir binding, which would most likely prevent Sas2 from binding) or methylated H3-H4 dimers.
- **Dot1 methylation**. Dot1 is a methyltransferase that methylates H3K79. Its catalytic rate is greatly enhanced in H3-H4 dimers which are already H4K16-acetylated (and H2B ubiquitilated [2]). Hence, we assume for simplicity that it can only methylate those H3-H4 pairs with the H4K16ac mark. We model the methylation reaction at the locus with a propensity function linear with the number of acetylated H3-H4 histone pairs:

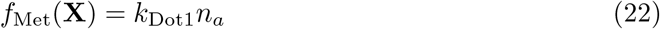
- **Sir2 deacetylation**. Sir2 is the catalytic subunit of the SIR complex, which deacetylates H4K16. We model the deacetylation reaction with a propensity function proportional to the product of Sir complexes at the locus and acetylated H3-H4 dimers (whether methylated or not):

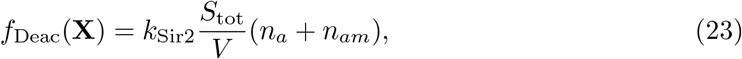

where *S*_tot_ is the total number of Sir complexes bound to the locus, including the silencers: *S*_tot_ = 2(*n*_*en*_ + *n*_*in*_ + *n*_*ei*_) + *n*_*d*_. Being a two-particle reaction, its propensity is divided by the volume of the locus *V*, since a larger volume would dilute the concentration of Sir-bound complexes and acetylated H3-H4 histone pairs, thereby reducing the number of contacts between them and the rate of the deacetylation reaction.

These propensity functions are used by the simulation in order to compute how likely it is that a given reaction will occur within a (small) time frame.

#### 1.4 Cell cycle: chromosome replication

Finally, upon DNA replication, each nucleosomes is lost with probability 0.5 and, if so, replaced by an acetylated nucleosome. Given that in the model we consider each H3-H4 pair as independent, every time a nucleosome is replaced, two H3-H4 histone pairs (H2A and H2B are irrelevant for the model) are chosen at random between the remaining (non-replicated) nucleosomes for replacement. In addition, at replication, we assume all Sir proteins bound at the locus are lost, independently of whether the nucleosome is replaced or not. Given that the fastest timescale in the system is Sir protein binding, we expect to recover them shortly after replication in most cases, but not all. For a detailed explanation of the implementation of this step, see Section 2.2

#### 1.5 On the volume of the locus

The model as described above (assuming *V* constant) does not display a clearly bistable behaviour. This is partially due to the fact that, unlike in previous theoretical studies [6], here we do not have long-range cooperativity between Sir proteins (we restrict this cooperativity to the dimerization reaction) and, thus, the only source of cooperative silencing is the feedback between Sir2 deacetylation and Sir binding (which, apparently, is not strong enough). We therefore needed a *stronger non-linearity* that makes the expressed and silenced states more polarised.

In this section we develop the hypothesis that the additional ingredient that stabilises the two distinct states is the *varying volume of the locus*. If the locus shrinks upon SIR binding or it expands when nucleosomes are acetylated or methylated, the states become more polarised and the transitions are less frequent. This is because a smaller locus would favour two-particle interactions, which happen to be the silencing-establishing ones (Sir binding and Sir2 deacetylation). More precisely, a small locus favours deacetylation and Sir binding, which in turn maintains the locus small, while a expanded conformation of the locus would inhibit them, effectively favouring acetylation and methylation and keeping the locus expanded. Thus, even if there is no explicit long-range cooperativity, the changes in locus volume act as a sort of locus-wide cooperative behaviour.

There have been a number of observations and simulations that suggest views similar to this one. In Ref. [9], chromosome conformation capture experiments at HMR imply that the locus is compacted in the wild type but more expanded in a *sir3 Δ* mutant. It has also been reported that chromatin acetylated at H4K16 presents a more open configuration, as opposed to the compact 30nm fiber [10], and that Sir2 deacetylation of chromatin *in vitro* makes it more compact [11]. In addition, recent computational simulations suggest that chromatin carrying the H4K16ac mark acquires an expanded conformation, driven by interactions including the disordered H4 tail [12], and, in particular, H4K16 - DNA contacts [13].

Then, based on these observations, we assumed that the volume of the locus depends on H4K16ac and, mathematically, is given by

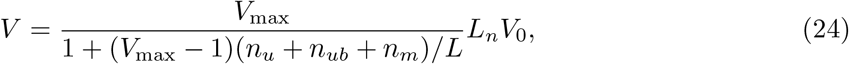

where *n*_*ub*_ = *n*_*d*_ + *n*_*en*_ + *n*_*in*_ is the total number of Sir-bound H3-H4 pairs, *L*_*n*_ is the number of nucleosomes, *V*_max_ is the ratio between the maximum and the minimum volume of the locus, in such a way that *V* ∈ [*L*_*n*_ *V*_0_, *L*_*n*_ *V*_0_ *V*_max_], and *V*_0_ is the unit volume we consider. In this case, *V*_0_ was taken to be the volume occupied by a nucleosome. Finally, the fact that *V* ∼ (*n*_*u*_ +*n*_*ub*_ +*n*_*m*_)^−1^ reflects the dependence of locus size with deacetylation of the H4 tail, since *n*_*u*_ + *n*_*ub*_ + *n*_*m*_ is the number of histone pairs that are not acetylated. For a simplified polymer model justifying this functional form, see Section 3.

Eq. (24) is used in Eq. (23) and in the Sir dimerization rates (Section 1.1), reducing the rates of deacetylation and dimerization as the volume increases.

#### 1.6 Parameters

The model parameters used in the rest of this document take the following values, unless otherwise specified:

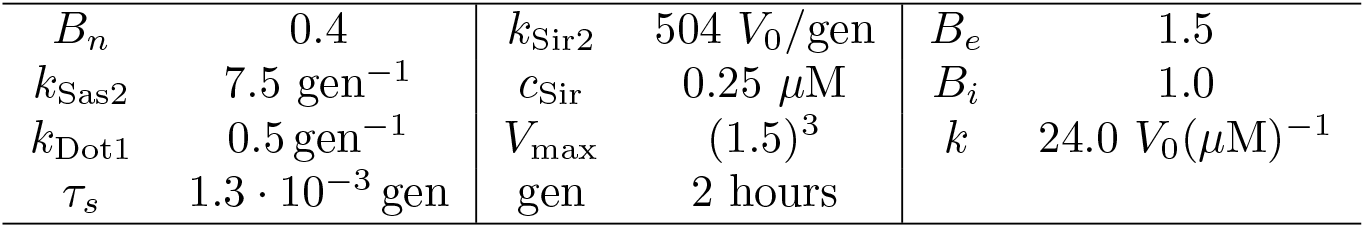

Time is given in units of a cell cycle length, taken to be two hours. *V*_0_ is the unit volume, which is the characteristic volume of the system; in this case, roughly the volume occupied by a nucleosome. Note that *V*_0_ only appears in rate constants that are divided by the volume to compute the rate, thus creating a rate which has units of the inverse of time and is independent of *V*_0_.

Moreover, we choose *k*_Dot1_ ≪ *k*_Sas2_ because we are considering a histone deacetylase (Sir2) but not a demethylase, and thus the only way to dilute the methylation mark is by replication, which is the slowest timescale in the system. On the other hand, the fastest timescale in the system is *τ*_*s*_, which corresponds to the Sir binding and unbinding dynamics. Its value corresponds to roughly 10s in the units of the yeast cell cycle [3].

When modelling mutant strains some of these parameters will be varied. For example, in the silencer strength mutant *sir*1*Δ*, we take *B*_*e*_ = *B*_*i*_ = 0.6 to mimic the weakening of silencers (in this mutant the enrichment seems to be similar at the silencers and across the locus). If comparing to data obtained from strains that bore a tag for immunoprecipitation (myc or V5) *k* is reduced to 8.4*V*_0_(*μ*M)^−1^, representing a lower binding affinity due to the tag. In the case of the V5 tag, *B*_*e*_ is also reduced to 1. Finally, we take *c*_Sir_ = 0.25 *μ*M (half of the Sir3 concentration estimated in from Refs. [3, 6]), although this parameter will be varied in the Sir titration.

These were the parameters we used for most of the study after correcting for the steep trend in the experimental silencing establishment rate. The original parameters put less weight on the silencers and, thus, predicted a milder trend. The results of the original parameters can be seen in Fig. 2A,B in the main text and Supplementary Figure 3 (below). The original parameters for the *sir*1*Δ* mutant differ from the final ones only in the values of *k* = 12.5*V*_0_(*μ*M)^−1^ and *B*_*e*_ = *B*_*i*_ = *B*_*n*_ = 0.5.

### 2 Computational simulation of the system

In Section 1 the model has been described in detail and in this section we specify the mathematical and computational framework to simulate such a model. The dynamics are divided in two separate parts: the dynamics during the cell cycle (protein binding and histone PTMs) and replication. The Fortran implementation of this simulation can be found at: https://github.com/AMovillaMiangolarra/Conformation Acetylation Feedback 2023

### 2.1 Cell cycle dynamics

The processes taking place during the cell cycle have been specified in Sections 1.1 and 1.3. Given these processes, the evolution of the probability of the system being at state **X** is given by the chemical master equation

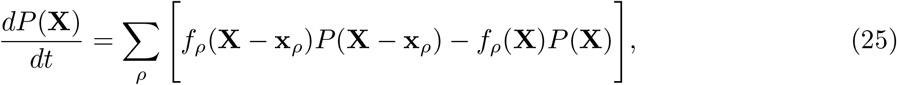

where the index *ρ* runs over all the processes specified in Sections 1.1 and 1.3. *f*_*ρ*_(**X**) specifies the rate at which a process *ρ* occurs when the system is at state **X**, whose functional form was specified in Sections 1.1 and 1.3. Finally, **x**_*ρ*_ is the vector that encodes for changes in the system due to the process *ρ*. For example, in a methylation reaction, **x**_*ρ*_ is a vector that subtracts an acetylated H3-H4 pair from the system (**X**) and adds an acetylated and methylated one. For further details and a more complete exposition of the chemical master equation, we refer the reader to one of the reviews or books written on the subject, for example [7].

In order to simulate stochastic trajectories that satisfy Equation (25) we used the Gillespie algorithm, which operates in continuous time by taking advantage of the exponential waiting times between subsequent reactions. The method is explained in detail in Ref. [14]. Briefly, in each iteration, we obtain two probability distributions: one for the time at which the next reaction will occur and another one for the type of reaction (whether it will be Sir binding, acetylation, methylation or deacetylation). Two random numbers are then drawn to sample these distributions stochastically.

### 2.2 Replication

At the end of a cell cycle the chromosome is replicated and each nucleosome is randomly inherited from the mother strand to one of the two daughters. This means that there will also be some incorporation of new nucleosomes into the chromatin in order to fully repopulate the daughter strands.

The process to simulate the inheritance of histone pairs is as follows:

1. Start with a histone pair pool equal to the state **X** but with the bound Sir proteins removed (and ignoring silencer states).
2. Obtain a random number in *r* ∈ [0, 1), uniformly distributed.
3. Obtain another random number *r*_1_ ∈ [0, 1), uniformly distributed that specifies the first histone pair. Histone pairs are ordered and are selected if *r*_1_ falls within their assigned interval of width the inverse of the number of histone pairs remaining in the pool.
4. Remove the chosen histone pair from the histone pair pool.
5. Repeat steps 3 and 4, with another random number *r*_2_, to choose the second H3-H4 pair in the nucleosome.
6. If *r <* 0.5, the nucleosome (including the two histone pairs) is lost and is replaced in the daughter strand by two H3-H4 pairs that are acetylated but not methylated. If *r* ≥ 0.5, then the nucleosome is inherited by the daughter strand.
7. Repeat steps 2-6 until the histone pair pool has been depleted.

At that point the whole locus will have been replicated and the simulation will correspond to the state of chromatin at the locus of one of the daughter cells.

### 3 Scaling proposal for locus volume

In Eq. (24) we state a relation between locus volume and the acetylation status of the chromatin. Here, we seek to justify this assumption under several approximations, within the simple framework of the Flory mean-field theory for polymers [15].

In the silenced state, the locus is compact and highly deacetylated. In this limit, we can consider the nucleoplasm a poor solvent for the locus (due to, mainly, the attractive interactions between deacetylated H4 tails), where an interplay between attractive and repulsive (mostly steric or excluded volume) interactions are the strongest determinants for chain conformation. For highly compact chains the entropic contributions to the free energy are negligible and, in the case of low acetylation, the effective two-body interaction is attractive. Then, the free energy of the system can be estimated by

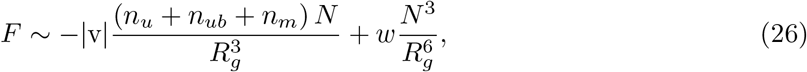

where *N* is the number of nucleosomes, *R*_*g*_ is the radius of gyration, v is the effective two-body interaction (v *<* 0), and *w* the three-body interaction [16]. The attractive two-body interaction is proportional to (*n*_*u*_ + *n*_*ub*_ + *n*_*m*_) *N* because unacetylated H4 tails are thought to drive compaction by contacts with DNA and other histone cores [12, 13]. However, in compact chains (a closely packaged locus), the three-body interaction is strong (last term in Eq. 26) and is dominated by excluded volume terms, which make *w* strongly positive and prevent an unphysical collapse of the chain.

If the chain is allowed to equilibrate with respect to its physical conformation, which should be the case in long timescales (minutes, hours), then the volume of the locus is given by the minimisation of the free energy, yielding

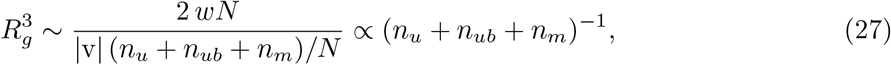

which is the scaling used in Eq. (24).

We emphasise that the scaling derived here is only valid in the limit where the density of nucleosomes is high enough (in order to neglect entropic terms) and the two-body interaction is negative and dominated by deacetylated H4 tails. This approximation, together with Eq. (24), might not be accurate for expanded and active loci.

More accurate expressions for the locus volume could be given for both active and silenced (expanded and compact) loci, but given the lack of distance and contact measurements at the scale of the HMR locus, we deemed this approximation sufficient for our purposes.

### 4 Modelling of mutant strains

There are several mutant strains considered throughout this work.

Silencer strength mutants are taken into account by decreasing the affinity of the SIR complex to the silencer. For *sir1Δ, B*_*e*_ = 0.6 and *B*_*i*_ = 0.6; while for *rap1 b*.*s*.*Δ, B*_*e*_ = 1.0 and *B*_*i*_ = 0.8.

Another category of mutants are those which affect the function of the SIR complex. The *sir3-bahΔ* mutant involves a deletion in the nucleosome binding domain. Hence, we model it by decreasing the affinity of the SIR complex to nucleosomes using *B*_*n*_ = 0.24. The other SIR complex mutant considered is the *sir2-N345A*, which makes Sir2 catalytically inactive. This mutant is modelled by setting the rate of Sir2 deacetylation to zero.

The other type of mutants considered are histone mutants, whose modelling is slightly more complex and is explained below.

#### 4.1 Modelling of *H4K16Q*

*H4K16Q* is supposed to mimic acetylation at H4K16 and it has been shown that it reduces the binding affinity for the SIR complex [17]. However, it does not seem to be a good mimic in the context of chromatin decompaction [18].

Therefore, in the model, we decrease *B*_*n*_ to 0.3 and assume that the volume of the locus does not change, since it can no longer be dynamically acetylated or deacetylated. The volume is thus fixed to *V* = 2 *LV*_0_, which is a lower value than the maximum for the wild type, reflecting the fact that Q is a bad mimic of acetylation regarding chromatin conformation. Finally, we assume that, even if H4K16 is not strictly acetylated, H4K16Q is a sufficiently good mimic to allow Dot1 to methylate H3K79 and, thus, any H3-H4 pair that is not Sir bound can be methylated. We have therefore reduced the possible states of the histone H3-H4 pairs to only three: Sir bound, unmodified (but low affinity due to H4K16Q) and H3K79me.

#### 4.2 Modelling of *H3K79L*

*H3K79L* is assumed to be a good methylation mimic. The possible states of histone H3-H4 pairs are reduced to three: Sir bound, unmodified (but low affinity due to H3K79L) and H4K16ac, where we set *B*_*n*_ to 0.24 for the unmodified H3K79L histone pair.

### 5 Comparing titration switching rates

#### 5.1 Ratio between wt and estradiol

In order to keep the modeling within a common framework we used switching rates from the native promoter as a scale factor for the switching rates obtained in the Sir4 titration (which included an estradiol inducible promoter). Thus, the switching rates obtained from the model were multiplied by the ratio between the switching rates in *sir*1*Δ* with the estradiol promoter (from [5]) and the native promoter (from [19]). Probably due to more noisy transcription in the estradiol-inducible promoter, this ratio is approximately 2.

#### 5.2 Titration values

As explained in Section 1.2, in order to compare with the output of the mathematical model, we link estradiol concentration with Sir complex concentration using a linear relationship. The parameters used were *α* = 1*/*1060 and *β* = 0.025 *μ*M.

### 6 Bimodality coefficient

The bimodality coefficient (BC) is a metric that quantifies the extent to which a probability distribution is bimodal. It takes values from 0 to 1, with higher values being more bimodal [20]. For example, for a Normal distribution the value is 1/3, for a uniform one ≃ 0.56 and for the sum of two different Dirac delta functions it takes its maximum value of 1 [21]. Values larger than 0.56 are often taken to correspond to bimodal distributions, which in the case of this work corresponds to bistable strains. Mathematically, the bimodality coefficient is given by

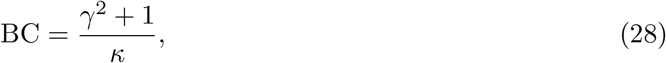

where *γ* is the skewness and *κ* the kurtosis of the distribution.

The BC for the model is obtained from the probability distribution for the fraction of histones acetylated in the locus. For the experimental data, the BC is computed from the Green Fluorescent Protein (GFP) levels for each condition (data from Ref. [5]). Given that the GFP levels were obtained in arbitrary intensity units (imaging experiments) and that this metric is heavily influenced by extreme values, we take the GFP levels in a logarithmic scale to minimise this bias and reduce the impact of noise.

### 7 Supplementary Figures

#### 7.1 Supplementary Figure 1: GFP levels with larger loci

As explained in the main text, in order to test the predictions of the model, strains with the *sir*1*Δ* mutation and larger HMR loci were created. To this end, different intergenic sources of DNA were inserted (HML, *K. lactis* Leu2, *K. lactis* Ura3) at different locations of HMR (R or L) depending on whether they were inserted at the right side or left side of HMR (see Supplementary Figure 1A). Only the strains whose added DNA source was HML were kept for the quantification of the switching rates, but based on these data all the strains behave similarly, since upon addition of DNA to the locus the percentage of cells expressing GFP increases, see panel B.

#### 7.2 Supplementary Figure 2: Switching rates in larger loci

The model predicted a decrease in the silencing establishment rate for strains with more nucleosomes in the locus but no or little decrease in the silencing loss rate. The experimental values for the switching rates of strains with 12 (the wild type), 14 and 16 nucleosomes at HMR are shown in Fig. 2. In line with the model predictions, the changes in silencing loss rates were not statistically significant, while most of the pairwise comparisons for the silencing establishment rates were statistically significant (*p <* 0.05).

#### 7.3 Supplementary Figure 3: Model results with the original parameter set

After obtaining the results for larger loci (14 and 16 nucleosomes at HMR) the model was reparameterised slightly. For completeness, in Supplementary Figure 3 we show the results corresponding to Figure 2 in the main text with the original parameter set. For details regarding the parameters, see Section 1.6.

**Supplementary Figure 1:**
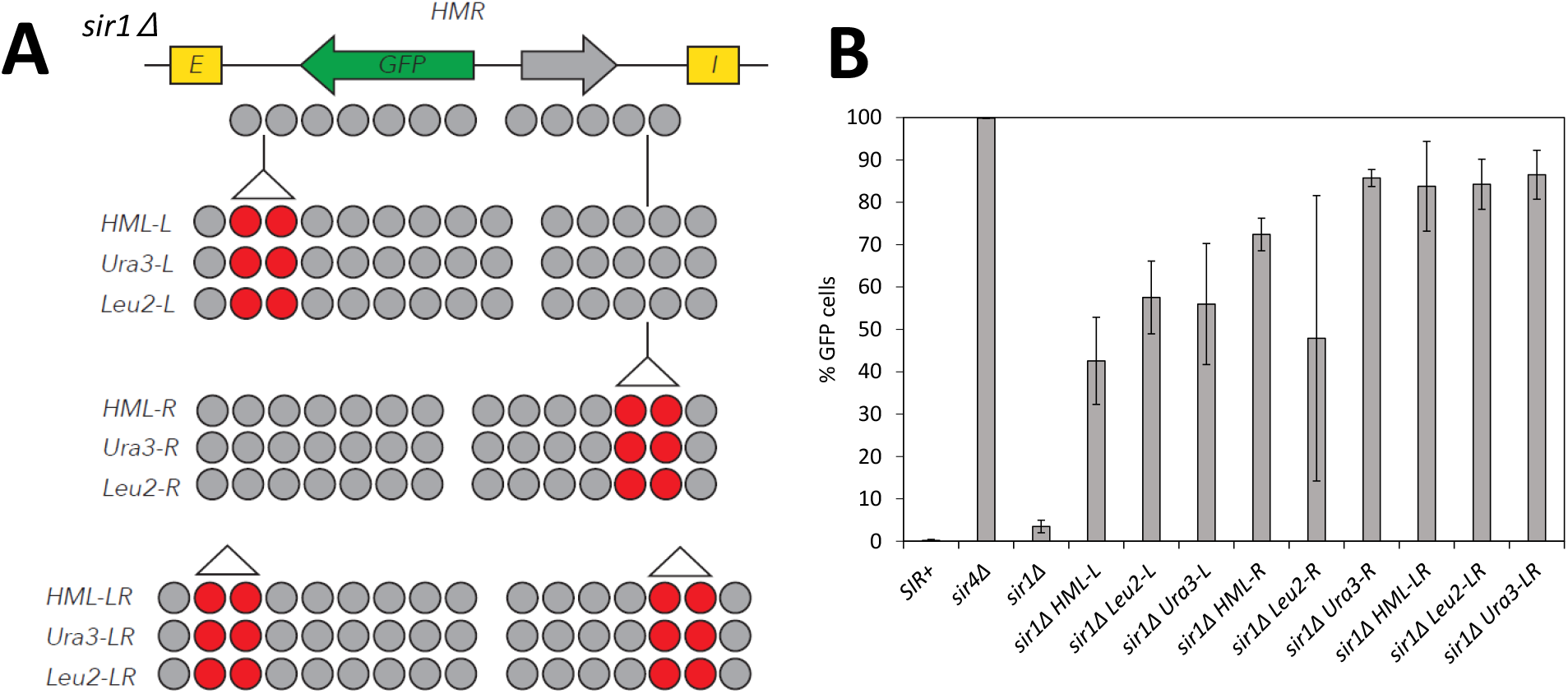
A) Schematic of the strategy for enlargement of the HMR locus. B) Percentage of cells expressing GFP at the HMR locus in different backgrounds. The error bars correspond to the standard deviation of 3 different replicates. *SIR+* is the control for a silent cell, since it includes the full SIR complex machinery to silence HMR, while *sir*4*Δ* is unable to silence the locus because it lacks Sir4 and thus nearly 100% of the cells express GFP. *sir*1*Δ* is the strain lacking Sir1 with the wild type HMR length, and the rest of the strains are identical except that different genomic regions have been inserted at HMR to make the locus larger. Strains whose name finishes in -L or -R have 14 nucleosomes at HMR and strains whose name finishes in -LR have 16.

**Supplementary Figure 2:**
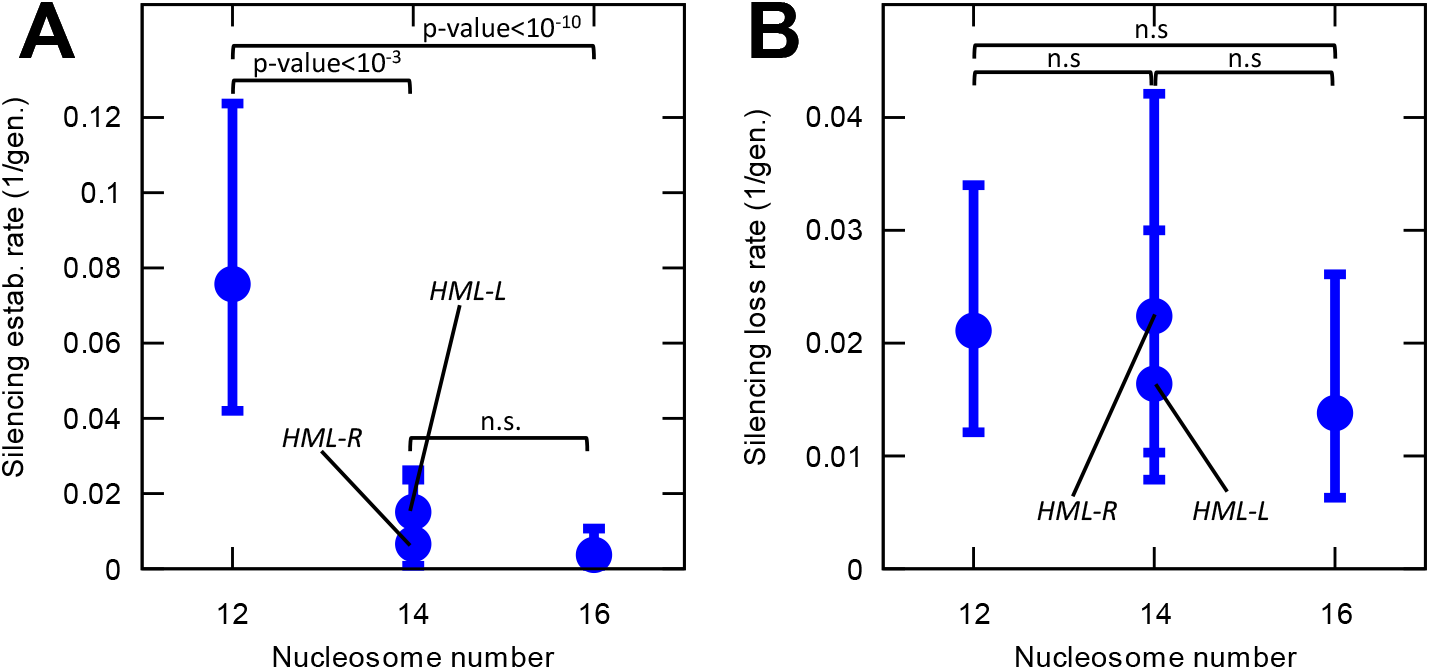
Experimental results for the switching rates as a function of nucleosome number. A) Silencing establishment rates for strains between 12 and 16 nucleosomes. B) Silencing loss rates for strains between 12 and 16 nucleosomes. There were two strains with 14 nucleosomes, since the intergenic HML DNA was introduced at one of two different locations to create a 14 nucleosomes locus. For 16 nucleosomes it had to be introducced at both locations and, thus, there was only one strain possible. For the statistical comparison, pairwise *χ*^2^ tests (with and without Yates correction) were performed and the most conservative p-value was taken. Not significant (n.s.) was taken to be a p-value over 0.05. The comparisons between *HML-L* and *HML-R* were not significant for both silencing establishment and silencing loss. The comparison between the silencing establishment of *HML-L* and *HML-LR* is significant (p-value*<* 0.05) but not so between *HML-R* and *HML-LR*. The error bars represent the 95% confidence interval, computed with the Clopper-Pearson or exact method.

**Supplementary Figure 3:**
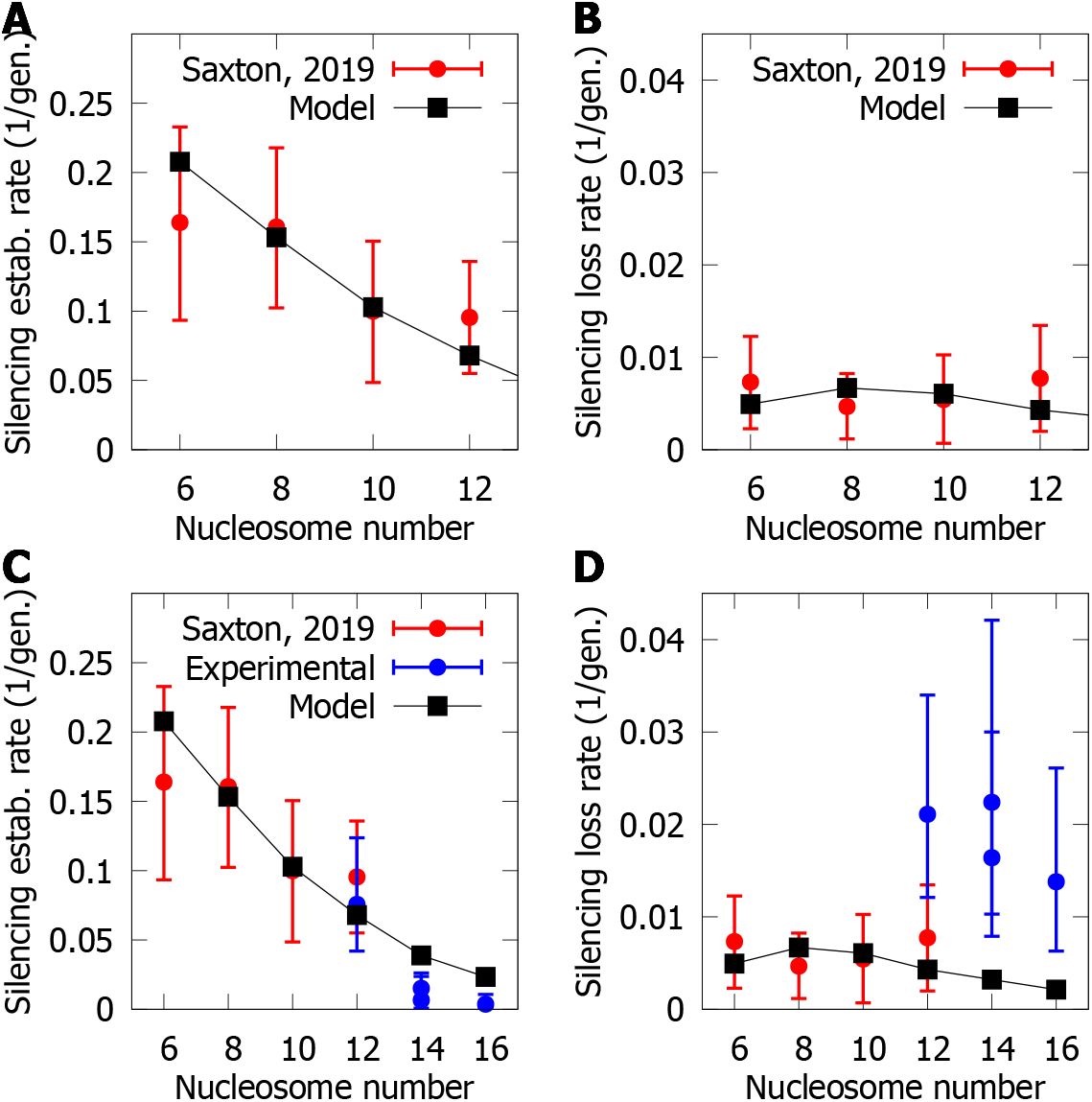
Switching rates as a function of nucleosome number. This figure is identical to Figure 2 in the main text, except that the results of the model in C and D were obtained with the original parameters.

#### 7.4 Supplementary Figure 4: Silencing establishment estimate

In experiments, by means of an estradiol-inducible promoter, it was possible to alter the concentration of Sir4 (and, thus, of the SIR complex) thereby modulating the switching rates of the strain. The modelling suggested that when increasing the concentration of the SIR complex, the silencing loss rate decays exponentially, but the silencing establishment does not increase exponentially. In Fig. 4, we show that the silencing establishment trend is not exponential with the Sir4 concentration either in experiments or simulations (see insets). Moreover, estimates of the silencing establishment rate based on fluctuations only at replication fail to reproduce the experimental or simulation data (see inset, green triangles), suggesting that other sources of fluctuations underpin the switch from expressed to silenced *HMR*. Supplementary Figure 4 is similar to Fig. 4A in the main text, except that the inset shows the silencing establishment rate and not the loss rate.

#### 7.5 Supplementary Figure 5: H4K16ac in the Sir4 titration

As explained above, with the estradiol-inducible promoter it was possible to lower the concentration of Sir4. Consequently, cells of the same strain were silenced when exposed to high estradiol concentrations, but expressed at low concentrations. We performed a similar procedure with the model, by varying the parameter *c*_SIR_ as explained in Section 1.2. To avoid having to model transcription and translation, we maintained the output of the model at the chromatin level. Thus, we obtained the overall level of H4K16 acetylation in the locus (which should correlate with transcription). The results for the wild type and two silencer strength mutants can be seen in Fig. 5.

**Supplementary Figure 4:**
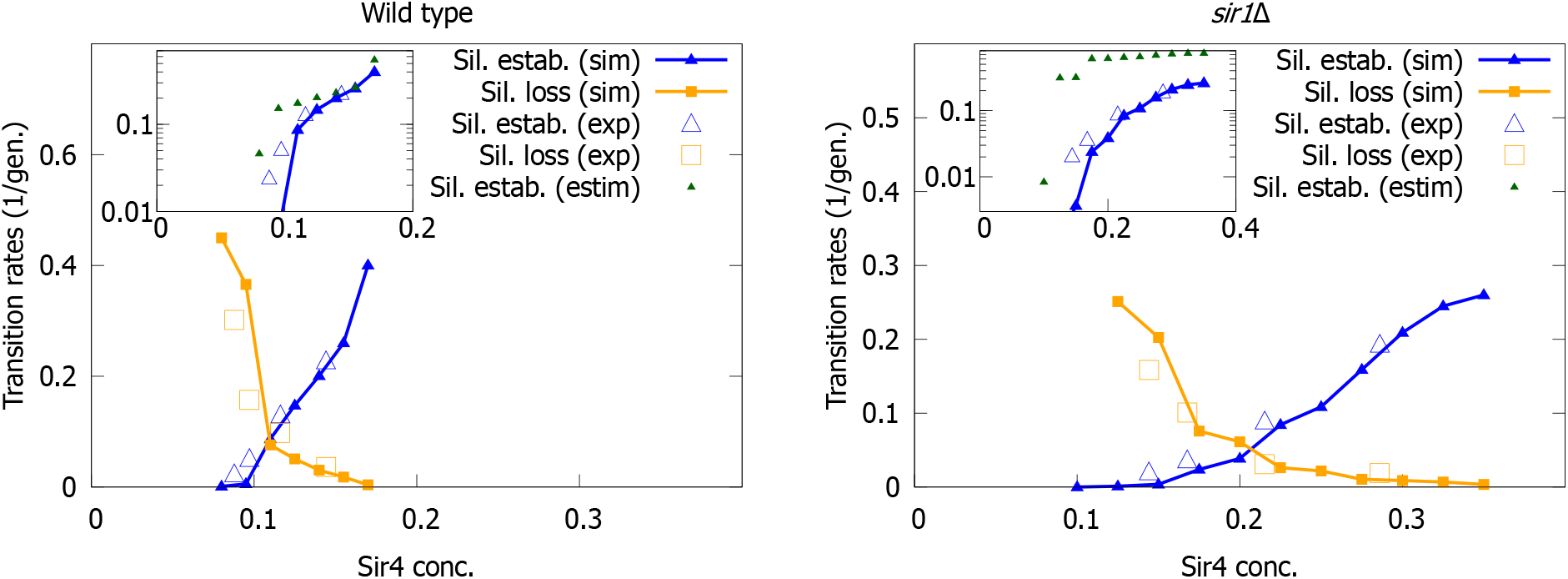
Switching rates in a Sir4 titration (identical to Fig. 4A in the main text except for the insets where, instead of the silencing loss, the silencing establishment rates are depicted). Silencing establishment and silencing loss rates in in the wild-type (left) and in a *sir1 Δ* background (right). Insets show the silencing establishment rates, in logarithmic scale, including an estimate based on the inheritance probabilities at chromosome replication, based on Eq. [2] in the main text. The experimental data was obtained from Ref. [5]. The poor agreement between the estimate and the full model (or the experiments) reflects the fact that fluctuations when distributing the nucleosomes after replication are not the main source of stochasticity for the switch from an expressed to a silent HMR.

**Supplementary Figure 5:**
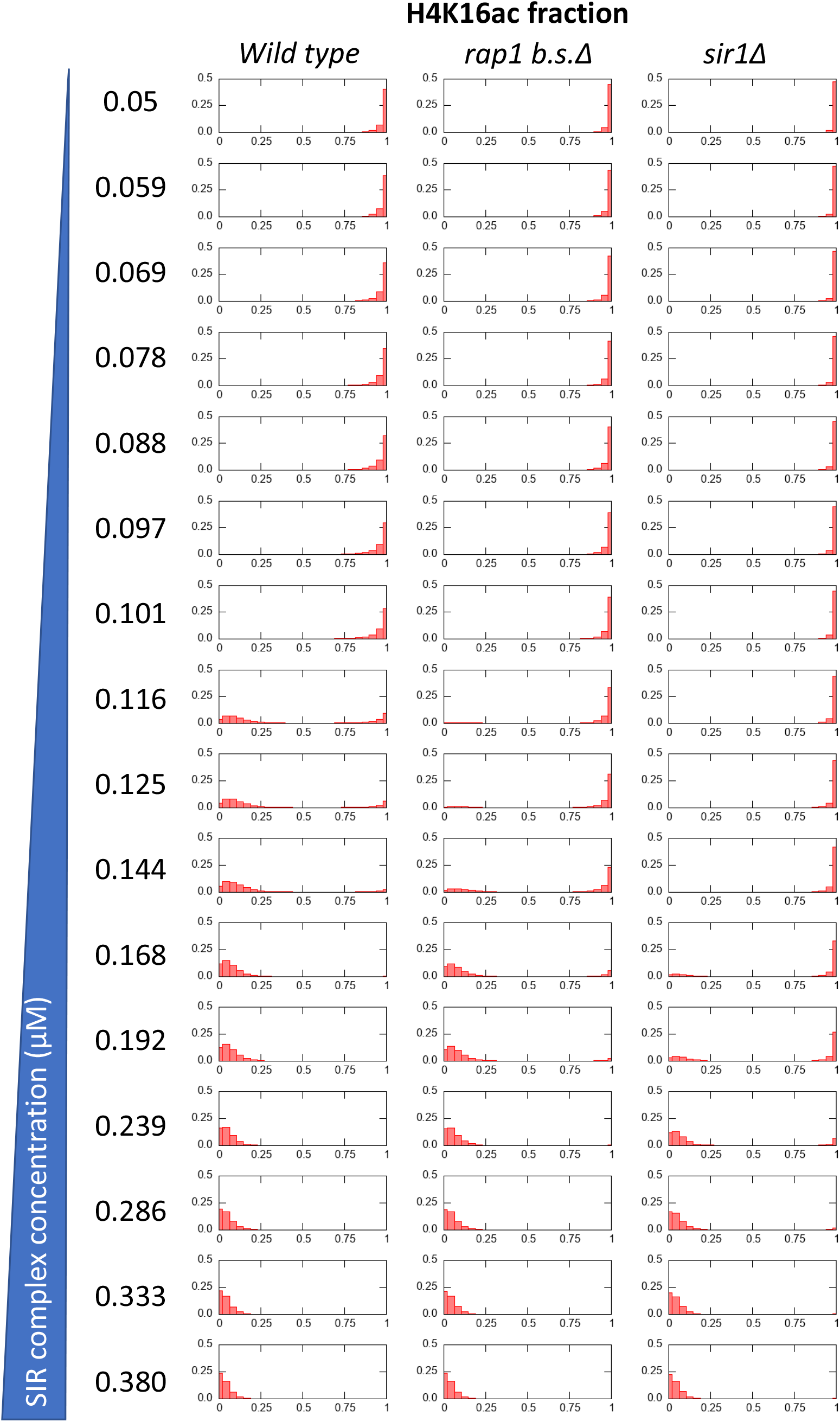
Probability of a given fraction of the HMR locus to be acetylated at H4K16 in three different silencer backgrounds (columns) and different SIR complex concentrations (rows). All three strains transition from an expressed state at low Sir concentrations to silenced at high Sir concentrations, passing through a concentration window in which they are bistable (two local maxima of probability). Importantly that concentration window is shifted towards higher Sir concentrations as the silencers are weakened.

---

**Table S1:**
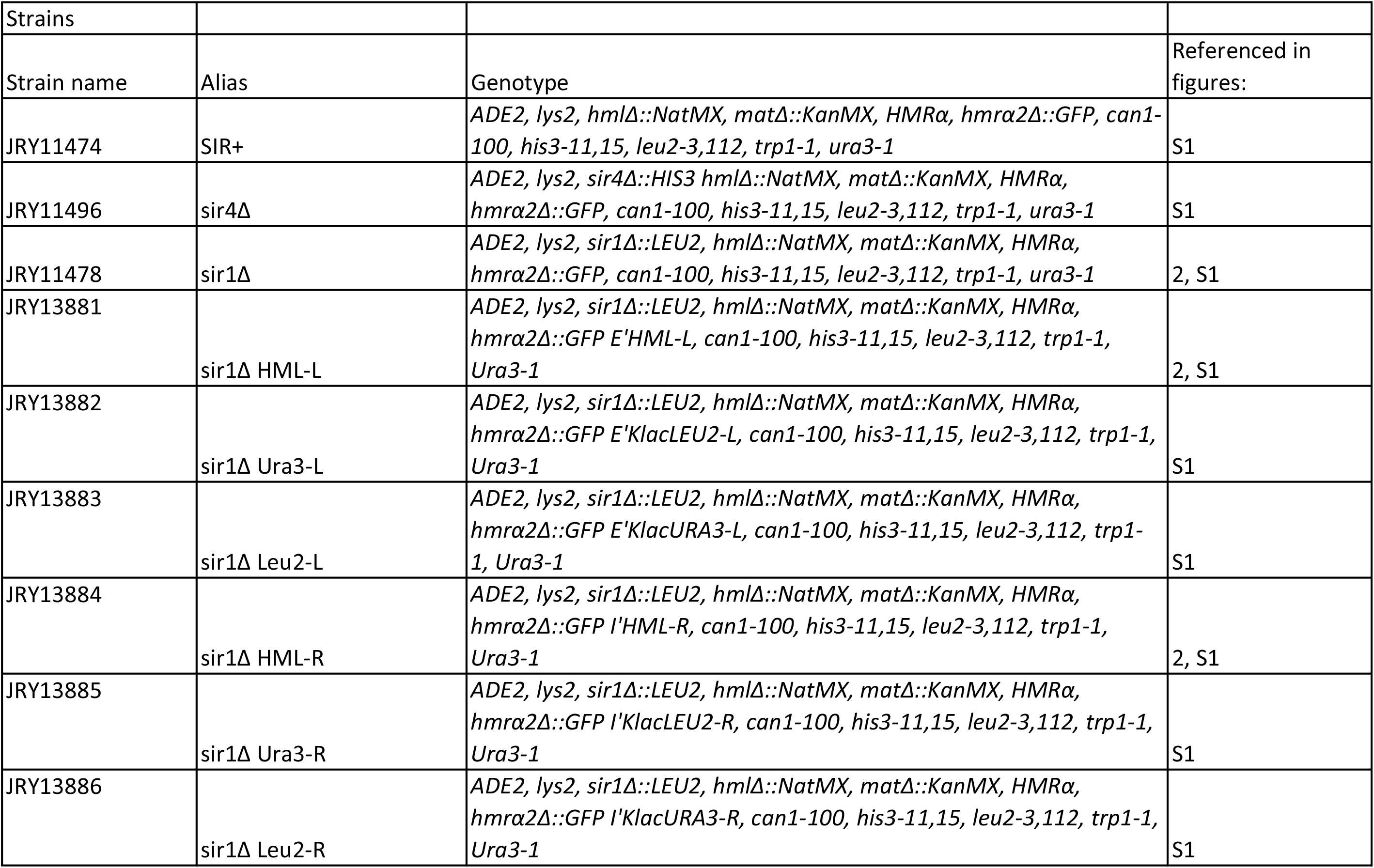
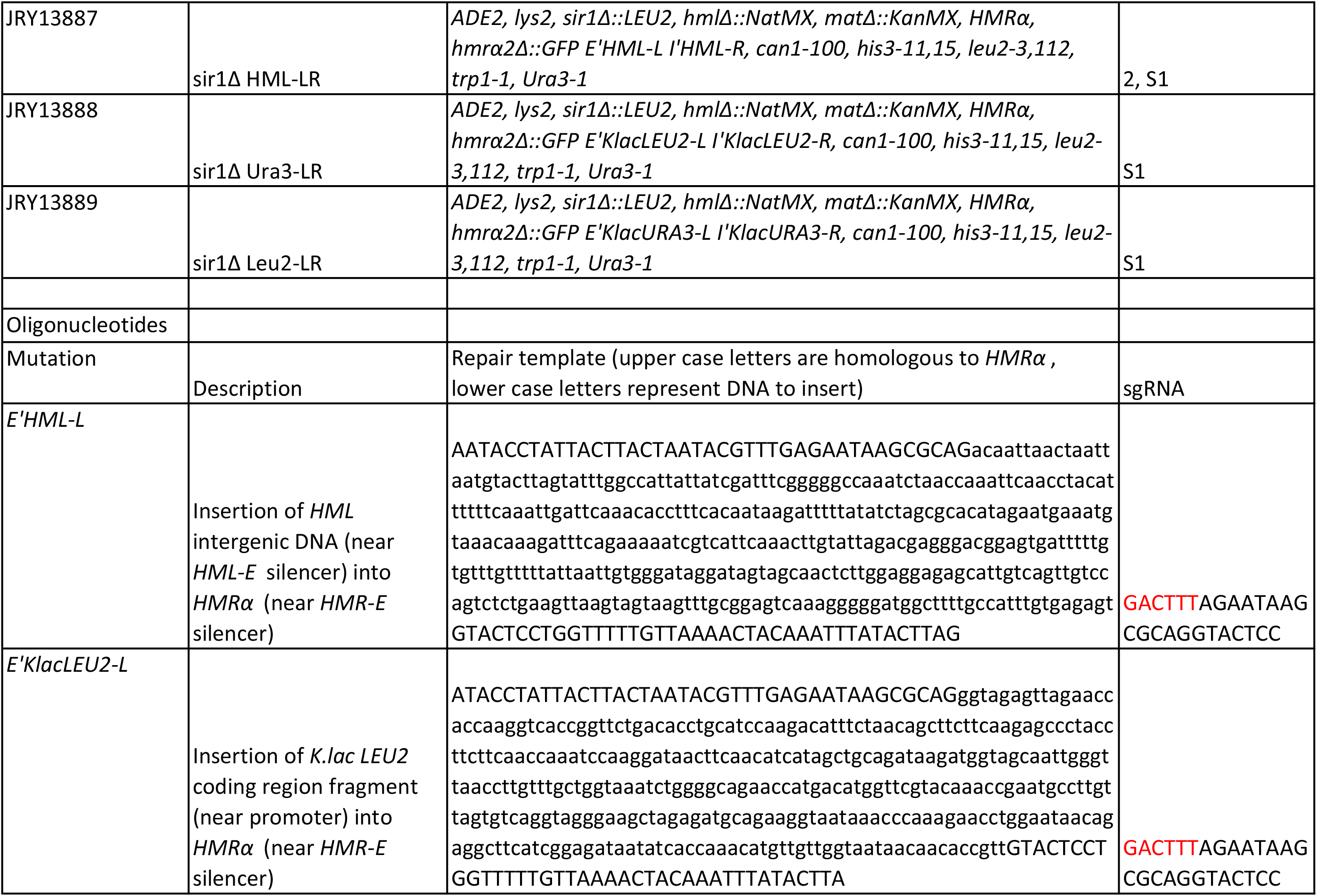
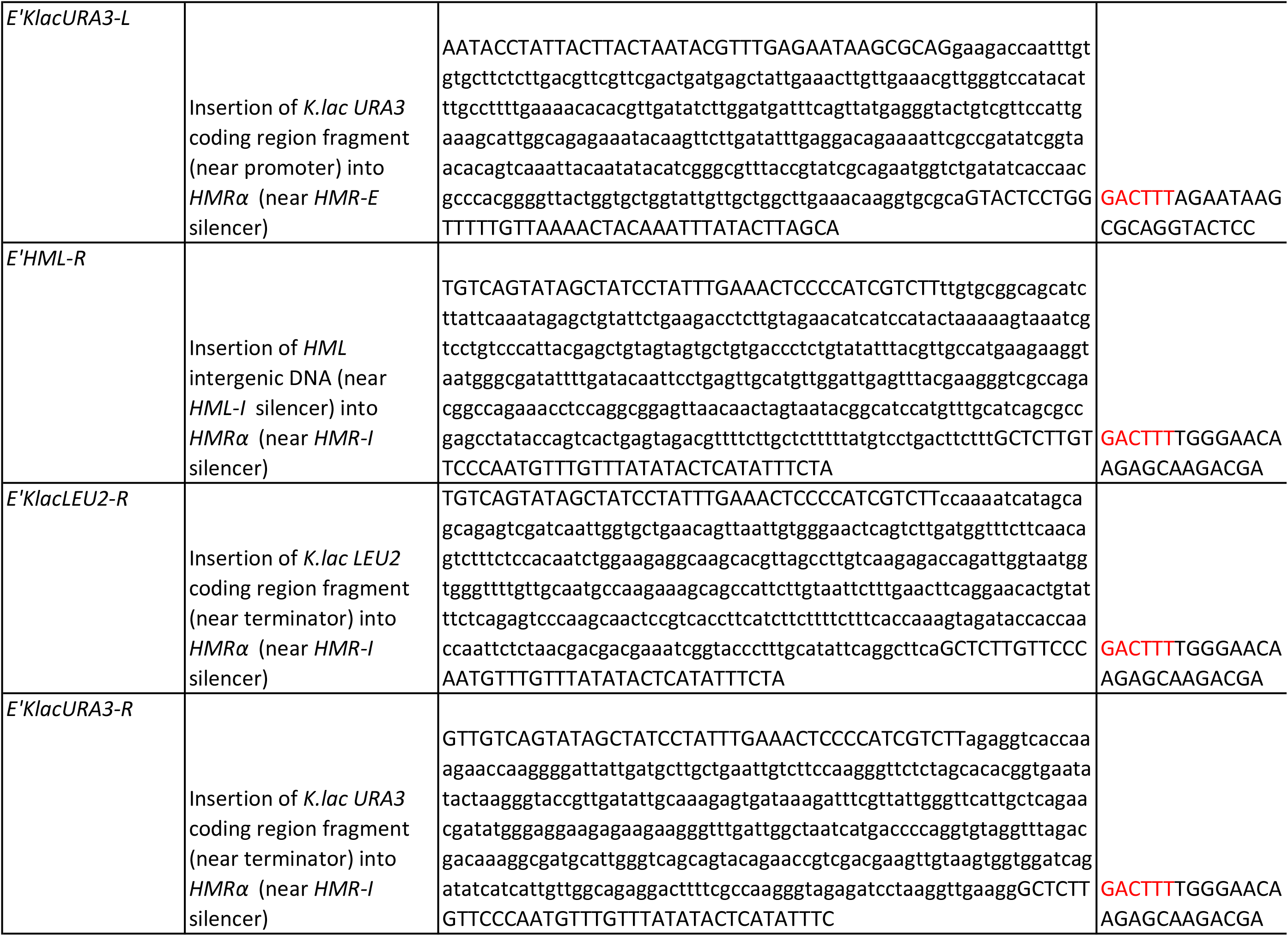
Strains and oligonucleotides

## Notes

https://github.com/AMovillaMiangolarra/Conformation_Acetylation_Feedback_2023

### Competing Interest Statement

The authors have declared no competing interest.

